# Reciprocal interaction between cortical SST and PV interneurons in top-down regulation of retinothalamic refinement

**DOI:** 10.1101/2025.01.18.633032

**Authors:** Qiufen Jiang, Sherry Jingjing Wu, Gord Fishell, Chinfei Chen

**Author notes:** Contact Information: Chinfei Chen F.M. Kirby Neurobiology Center Children’s Hospital Boston 3 Blackfan Circle Center for Life Sciences, CLS12250 Boston, MA 02115. Gordon Fishell Department of Neurobiology, Blavatnik Institute Harvard Medical School, Boston, MA 02115.

## Abstract

Refinement of thalamic circuits is crucial for the proper maturation of sensory circuits. In the visual system, this process is regulated by corticothalamic feedback during the experience-dependent phase of development. Yet the cortical circuits modulating this feedback remain elusive. Here, we demonstrate opposing roles for cortical somatostatin (SST) and parvalbumin (PV) interneurons in shaping retinogeniculate connectivity during the thalamic sensitive period (P20-30). Early in the refinement process, SST interneurons promote the strengthening and pruning of retinal inputs in the thalamus, as evidenced by disrupted synaptic refinement following their ablation. In contrast, PV interneurons, which mature later, act as a brake on this refinement, with their ablation leading to enhanced pruning of retinogeniculate connections. Notably, manipulating the relative balance between these inhibitory circuits can regulate sensory deprivation-induced retinogeniculate remodeling. Taken together, our findings show that cortical SST and PV interneuron circuits drive reciprocal antagonism that gate experience-dependent feedforward thalamic refinement.

## Introduction

Multiple aspects of the visual system are substantially reorganized as maturation proceeds. A key example of this is seen at the mouse retinogeniculate synapse, where the establishment of developmental circuits is followed by a second phase of refinement that is driven by sensory experience. ^1,2^ During this developmental window, called the thalamic sensitive period, external stimuli provide the signals for fine-tuning this connection between the eye and the visual thalamus. ^1–4^ For instance, visual deprivation during this period, referred to as late dark rearing (LDR), leads to an increase in the number of converging retinal inputs and a decrease in the average synaptic strength of the retinogeniculate connections. ^1,5^ In mice, this remodeling occurs between postnatal day (P)20 to P30, a process traditionally considered to be solely driven by bottom-up visual experience. Despite this canonical view, developmental studies of the visual system indicate that information during this period is far from unidirectional and that the primary visual cortex (V1) is both activated and centrally involved in bidirectional communication with the thalamus. ^6,7^ Multiple lines of evidence indicate that this top-down signaling plays a key role in directing the reorganization of retinogeniculate synapses. ^5,7,8^ Specifically, changing the activity of cortical feedback signaling alters retinogeniculate refinement. Increasing or decreasing the activity of layer (L) 6 corticothalamic neurons in V1 alters retinogeniculate connectivity during the thalamic sensitive period, ^5^ highlighting the importance of cortical-thalamic feedback in subcortical circuit refinement. ^9,10^ Given the need for precise coordination of cortical and thalamic maturation, a key question is how these processes are jointly regulated. How does visual experience processed within the cortex dynamically modulate top-down feedback to drive the remodeling of retinogeniculate synapses (Fig. 1A)? The precise timing during which the retinogeniculate circuits are reorganized provide a tantalizing hint that top-down corticothalamic signals regulating this process must also be precisely controlled. As infragranular pyramidal neurons (PNs) in V1 can be significantly tuned by local interneurons, ^11–14^ we speculated whether the maturation of inhibitory circuits regulating corticothalamic cells could play a central role in coordinating cortical and thalamic maturation.

**Figure 1.**
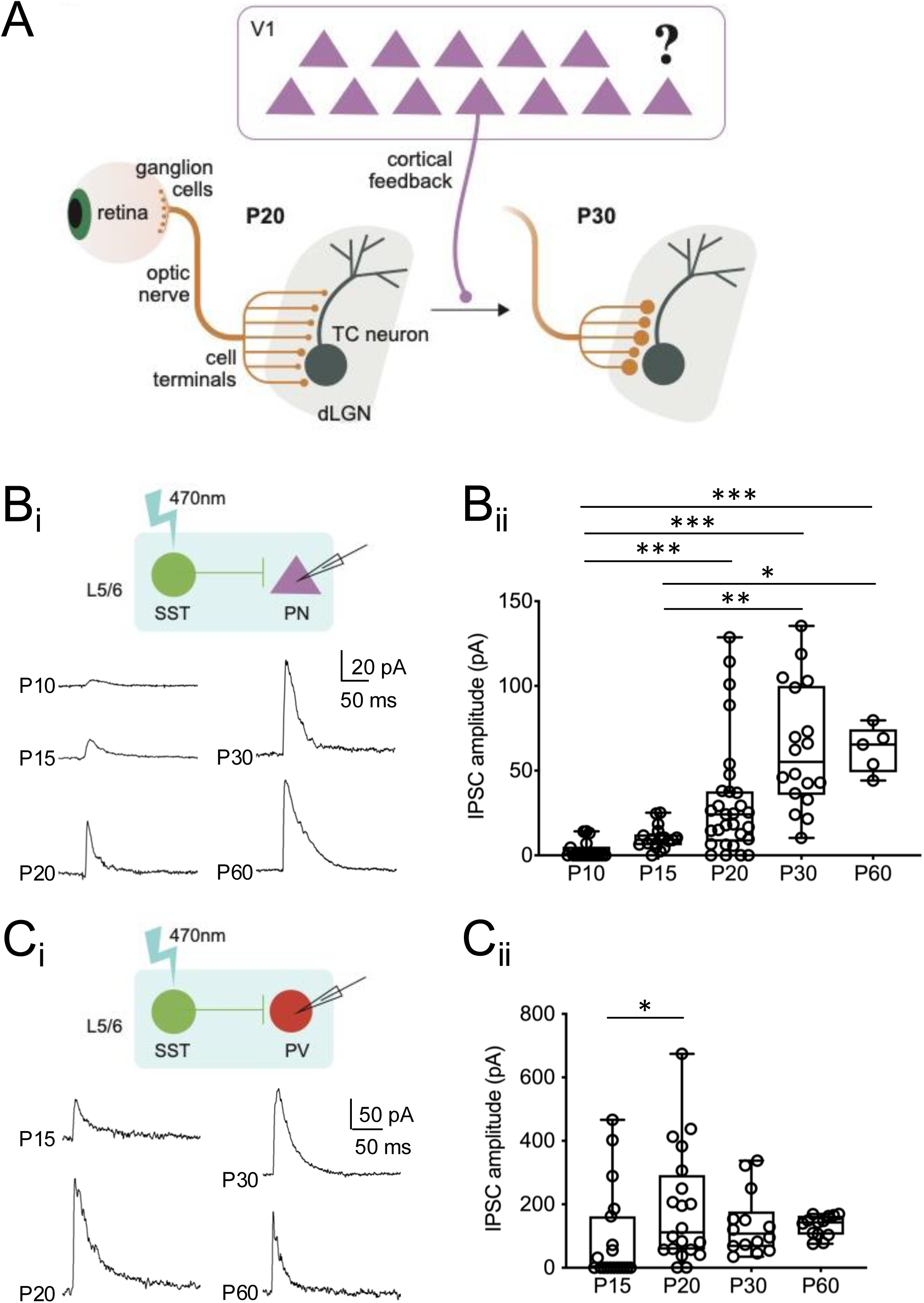
Maturation of somatostatin (SST) interneuron-mediated cortical circuits during the sensitive period for thalamic plasticity. A. Corticothalamic neurons in V1 L6 project to the dLGN and regulate the feedforward retinogeniculate refinement. ^5^ B. Developmental time course of inhibitory post-synaptic currents (IPSCs) from SST interneurons to pyramidal neurons (PNs) (SST→PNs) in V1 L5/6. B_i_ *Top*: schematic of optogenetic stimulation; *Bottom*: Example traces; B_ii_: Amplitude of SST→PN IPSCs over development. C. Similar to B but for SST→Parvalbumin (PV) interneuron IPSCs. B-C: * *P* < 0.05, ** *P* < 0.01, *** *P* < 0.001, Kruskal-Wallis test, Dunn’s multiple comparisons test.

Somatostatin (SST) and parvalbumin (PV) interneuron are two major interneuron types that modulate cortical function. The developmental regulation of these two populations is critical to the shift in cortical circuits from synchronous to decorrelated activity. ^13,15–23^ These developmental changes likely have a top-down effect on thalamic function regulated through corticothalamic feedback. To explore this idea, the output from SST interneurons onto PV interneurons and PNs in infragranular layers was examined in V1 using slice electrophysiology across development. Notably, the development of infragranular inhibitory circuits showed a similar sensitivity to sensory experience as retinogeniculate synapses within the same developmental time window, raising the possibility that the maturation of these inhibitory circuits directly drives the refinement of retinogeniculate synapses in the dorsal lateral geniculate nucleus (dLGN). To test this hypothesis, we manipulated the inhibitory circuits in V1 by ablating cortical SST or PV interneurons and evaluated the impact on retinogeniculate synapse refinement. Ablation of SST interneurons during the thalamic sensitive period disrupted both PV interneuron firing maturation and retinogeniculate synapse refinement. In contrast, ablation of PV interneurons during the same period enhanced retinogeniculate pruning. Moreover, activation of SST interneurons or ablation of PV interneurons during the sensitive period prevented the rewiring of retinogeniculate connections in response to visual deprivation. Taken together, our findings support a model in which SST and PV interneurons provide reciprocal antagonism to fine-tune the refinement of feedforward thalamic circuits. Our study highlights local interneurons as key regulators of experience-dependent interactions between cortical and thalamic circuits.

## Results

### Connectivity of SST interneuron circuits matures before the end of thalamic sensitive period

Infragranular PNs are a major source of cortical output. In L6 of V1, PNs send abundant long-range projections to first-order thalamus ^5,24–28^ and receive local inhibitory inputs from deep-layer interneurons. ^14^ Both SST and PV interneurons initially appear in infragranular layers during development (Fig. S1A-B) and thus are well positioned to modulate cortical feedback circuits. During the thalamic sensitive period, cortical inhibitory circuits are still undergoing developmental changes. To understand how these developing circuits impact corticothalamic feedback, we examined the functional responses of SST and PV interneuron circuits in infragranular layers of V1 over the developmental period from P10-60, spanning eye opening at ∼P14 and the thalamic sensitive period (P20-30). Due to the early onset of *Sst* expression, developing SST interneurons can be identified in V1 by immunolabeling at perinatal ages (e.g. embryonic day 20 in rat) ^29^ and can be genetically targeted using *Sst-IRES-Cre* line. When crossed to the *Ai14* reporter line, *Sst;tdTomato(tdT)* mice labeled a stable population of SST interneurons from P15-30 (Fig. S1A *top* and S1B *left*). In contrast, tracking the developing PV interneurons is more challenging due to the late onset of *Pvalb* expression. Although PV interneurons are present in the cortex earlier, ^30–32^ the expression of *Pvalb* turns on gradually between P10-30. ^33^ In *Pvalb-tdT* transgenic mice, only a small number of PV interneurons are genetically labeled before P20 (Fig. S1A *bottom*). Between P20 and P30 the number of labeled cells, confirmed by RNAscope *in situ* hybridization to be *Gad1*^+^ *Pvalb^+^* interneurons (see Methods, Fig. S1C-F), increases 22-fold (Fig. S1B *right*).

Since SST interneurons mature earlier than PV interneurons, we sought to characterize the developmental changes in the output circuitry of SST interneurons within infragranular layers. To achieve this, we recorded from PNs or genetically labeled PV interneurons (*Pvalb-tdT* mice) in L6 while optogenetically stimulating L5/6 SST interneurons using *Sst;ChR2* mice (progeny of *Sst-Cre* mice crossed with *Ai32*, a channelrhodopsin (ChR2) reporter line) (Fig. S2A-B, see Methods). PN neurons were recorded from L6 (see Methods) as we found that the vast majority of excitatory neurons in V1 L6 (∼84%) are labeled by Tle4, a marker for corticothalamic neurons (Fig. S1G-J). ^34–36^ Therefore, our PN recordings are enriched in corticothalamic neurons. The median peak amplitude of SST→PN inhibitory post-synaptic currents (IPSCs) increases steadily over an extended period starting from P10, reaching a plateau at P30 (Fig. 1B, Table S1). In contrast, the median strength of the SST→PV synapse increases significantly between P15 and P20, stabilizing thereafter (Fig. 1C, Table S2). The changes in SST synaptic charge transfer onto PN and PV interneurons over time, measured as the integral of the synaptic waveform, align with the observed amplitude changes (Fig. S2C-D). Therefore, by the end of the thalamic sensitive period, the infragranular inhibitory circuits mediated by SST interneurons have matured in V1, with the strengthening of SST→PV connection (P15-P20) occurring within the broader window of SST→PN development (P10-P30).

### Maturation of SST synaptic circuits in V1 is regulated by visual experience after eye-opening

Previous studies have shown that visual deprivation of mice between P20 and P30 (LDR), a significant change in sensory experience, triggers substantial rewiring of retinogeniculate connectivity, characterized by an increase in the number of retinal ganglion cell (RGC) inputs onto a thalamocortical (TC) neuron and a decrease in the average strength of these inputs. ^1,2^ If cortical interneurons actively participate in the process of experience-dependent retinogeniculate refinement by modulating corticothalamic feedback, we hypothesize that LDR may impact the development of these inhibitory circuits. To test this, we examined the inhibitory outputs from SST interneurons, including SST→PN and SST→PV synapses in L5/6 of V1 after LDR (Fig. 2), at a time point when inhibitory circuits have normally matured under standard light/dark conditions. The median amplitude of SST→PN IPSCs in LDR mice is significantly reduced (Fig. 2A_ii_ *left*, Table S1), and their cumulative probability curve is shifted to smaller amplitudes compared to controls reared in normal 12-hour light/dark cycles (NR, Fig. 2A_ii_ *right*). Notably, the strength of SST→PN synapses in LDR mice is reduced to a level comparable to that seen in P15 NR mice (*P* = 0.83, Mann-Whitney test), which is lower than the median strength observed at the onset of LDR in P20 NR mice (*P* < 0.05, Mann-Whitney test). In contrast, the amplitude of SST→PV IPSCs shows no significant changes after LDR (Fig. 2B, Table S2). Therefore, inhibition from infragranular SST interneuron to PNs is sensitive to sensory experience during the thalamic sensitive period.

**Figure 2.**
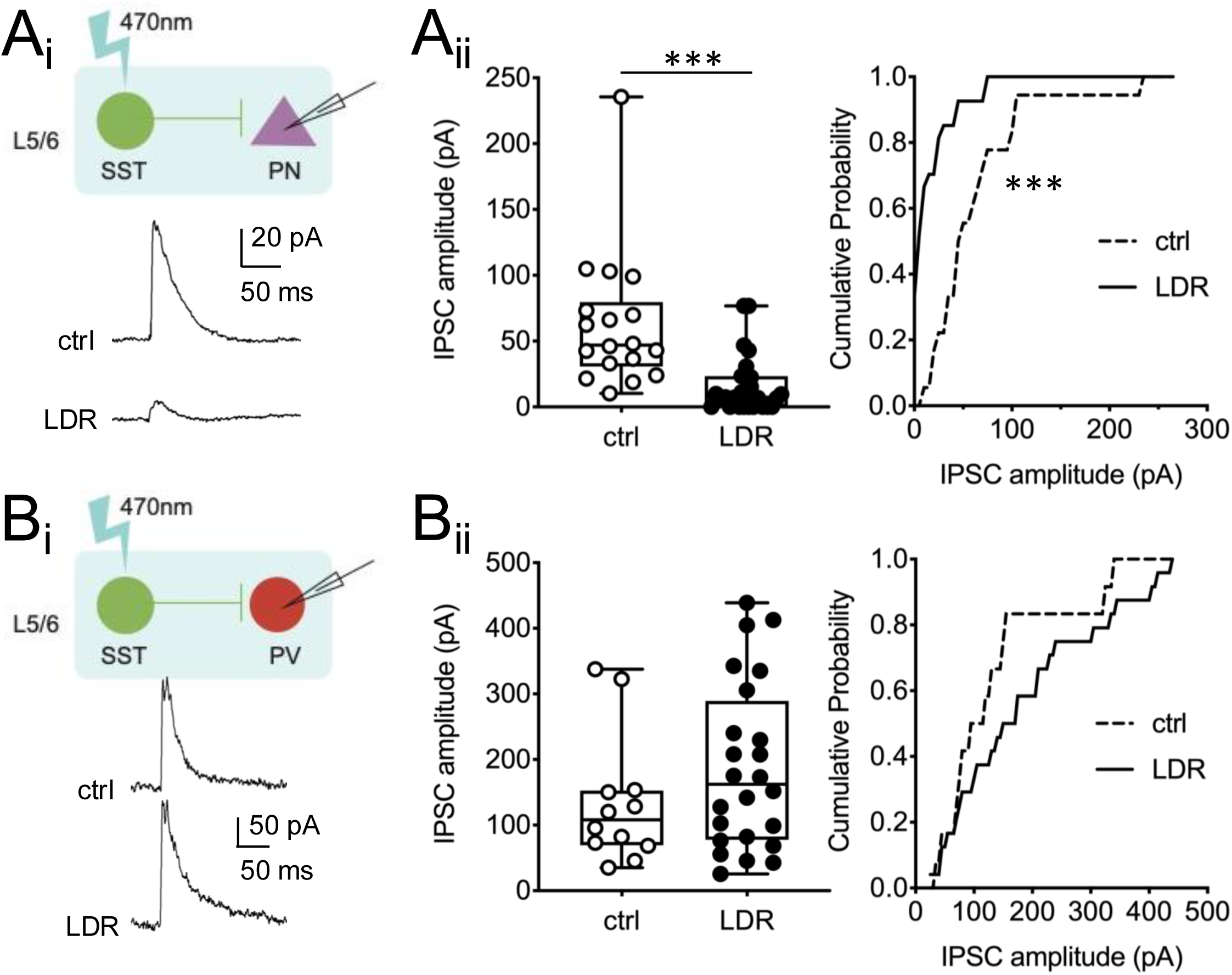
The maturation of SST interneuron-mediated circuits is regulated by late dark rearing (LDR). A. Effect of LDR on P30 SST→PN IPSCs. A_i_: Example traces; A_ii_ *left*: Peak amplitudes of SST→PN IPSCs, *** *P* < 0.001, Mann-Whitney test; A_ii_ *right*: Cumulative probability distribution of SST→PN IPSC amplitude, *** *P* < 0.001, Kolmogorov-Smirnov test. B. Effect of LDR on P30 SST→PV IPSCs. B_i_: Example traces; B_ii_ *left*: Peak amplitudes of SST→PV IPSCs; B_ii_ *right*: Cumulative probability distribution of SST→PV IPSC amplitude, Kolmogorov-Smirnov test.

### Ablation of SST interneurons during development disrupts pruning and strengthening of retinogeniculate synapses

Our findings that SST interneuron-mediated circuits mature concurrently with retinogeniculate refinement, and exhibit similar sensitivity to sensory experience, suggest a potential role for these cortical interneurons in thalamic circuit development. To test this hypothesis, we selectively ablated SST interneurons by injecting Cre-dependent Caspase3 (Casp3)-expressing virus into V1 of *Sst;tdT* or *Sst;ChR2* mice at P15 and assessed changes in retinogeniculate connectivity 15-20 days later (Fig. 3A). By P30, the majority of SST interneurons are eliminated across all cortical layers (Fig. 3B), resulting in a significant weakening of L5/6 SST→PN and SST→PV synapses (Fig. 3C-D, Table S1, 2). We then examined the effects of ablating SST interneurons on the refinement of retinogeniculate synapses by recording excitatory postsynaptic currents (EPSCs) from TC neurons in the dLGN using slice electrophysiology (Fig. 3E_i_). ^1,2^ Two parameters were used to assess synaptic refinement. First, single fiber (SF) strength quantifies the magnitude of the average retinal input onto a TC neuron. Second, the fiber fraction (FF), calculated as peak SF EPSC amplitude/maximum EPSC amplitude, measures the contribution of a single retinal input to total retinal drive, which estimates the degree of retinal input pruning. In normal development, retinogeniculate refinement leads to an increase in both FF and SF, where a higher FF indicates a reduction of converging retinal inputs due to pruning, while greater SF reflects an increase in synaptic strength. When compared to control mice, both the maximum retinal EPSCs and SF EPSCs are significantly weaker in mice expressing Casp3 in V1 SST interneurons (Fig. 3E_ii__-iii_, Table S3). The FF is also significantly reduced (Fig. 3E_iv_, Table S3), consistent with an increase in the number of convergent retinal inputs innervating each TC neuron. ^1^ Taken together, these results demonstrate that eliminating cortical SST interneurons leads to reduced retinogeniculate synaptic strength and pruning consistent with a less refined connection.

**Figure 3.**
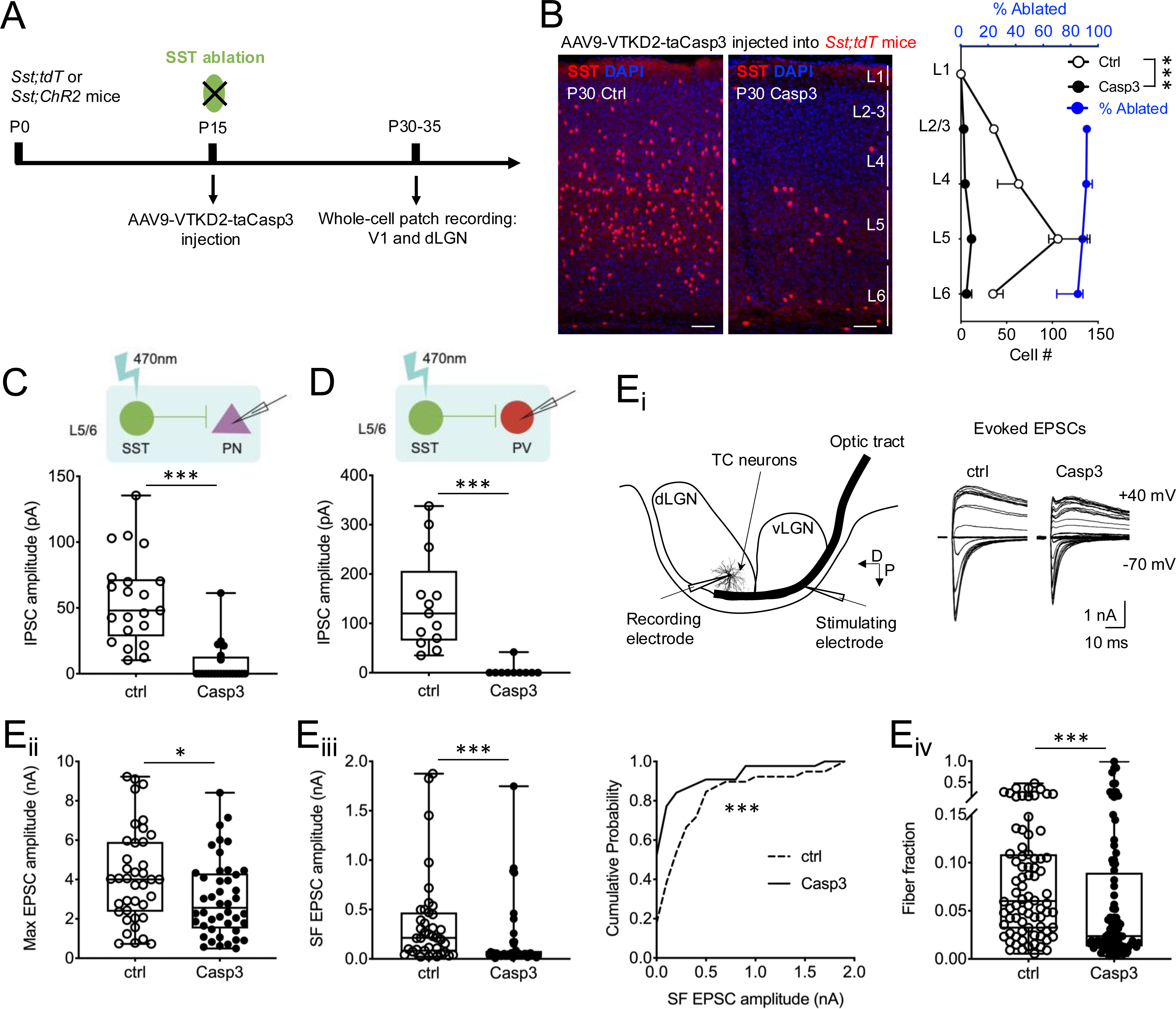
Ablation of SST interneurons during development disrupts normal retinogeniculate refinement. A. Protocol for ablating cortical SST interneurons. B. Ablation of cortical SST interneurons across different layers in V1. *Left*: representative images of SST-tdT^+^ interneurons in the V1 from mice injected with PBS (ctrl) or Casp3-expressing virus (Casp3). *Right*: Distribution of SST-tdT^+^ interneurons (black axis) and the proportion of SST interneurons ablated by Casp3 (blue axis) across V1, n = 8 mice. *** *P* < 0.001, two-way ANOVA. Scale bar: 100 μm. C, D. Effect of SST interneuron ablation on peak amplitude of (C) SST→PN IPSCs and (D) SST→PV IPSCs, both *** *P* < 0.001, Mann-Whitney test. E. Measurement of retinogeniculate responses after ablation of cortical SST interneurons. E_i_ *left*: Schematic of recording evoked excitatory post-synaptic currents (eEPSCs) from thalamocortical (TC) neurons in the dLGN by electrically stimulating the bulk of optic tract. D: dorsal; P: posterior; E_i_ *right*: Example overlaid traces of AMPAR and NMDAR-mediated eEPSCs by holding potential at -70 and +40 mV, respectively, in response to optic nerve stimulation; E_ii_: Peak amplitude of maximum (Max) AMPAR EPSCs, * *P* < 0.05, Mann-Whitney test; E_iii_ *left*: Peak amplitude of single fiber (SF) EPSCs, *** *P* < 0.001, Mann-Whitney test; E_iii_ *right*: Cumulative probability distribution of SF EPSCs, *** *P* < 0.001, Kolmogorov-Smirnov test; E_iv_: Distribution of fiber fraction (FF) ratio, *** *P* < 0.001, Mann-Whitney test.

### Ablation of PV interneurons during development enhances pruning of retinogeniculate synapses

Since SST interneurons also synapse onto PV interneurons, we investigated whether cortical PV interneurons also play a role in thalamic refinement. To directly assess their effects on retinogeniculate synapses, we used the same ablation strategy as for SST interneurons, deleting PV interneurons with *Pvalb-Cre* mice (Fig. 4A-B). Similar to SST interneuron ablation, the number of PV interneurons started to decline after 5 days of injection, and the majority were removed across different cortical layers by 15 days of injection (Fig. S4). We find that removing PV interneuron has a distinct effect on the pruning of retinogeniculate inputs onto TC neurons. The median amplitude of both maximum and SF retinogeniculate EPSCs does not differ significantly between PV-ablated mice and controls (Fig. 4C_i__-ii_, Table S3). However, the cumulative distribution plot of SF retinogeniculate EPSC amplitudes shows a rightward shift compared to controls, with a larger proportion of retinal inputs with strengths greater than 200 pA (Fig. 4C_ii_ *right*). Consequently, the FF significantly increases (Fig. 4C_iii_, Table S3), indicating fewer converging retinal inputs in the absence of cortical PV interneurons. Taken together, in contrast to SST interneuron ablation, which reduces pruning, ablation of PV interneurons enhances retinogeniculate pruning beyond normal levels by P30. These results demonstrate that while PV interneurons also regulate retinogeniculate synapse refinement, their role is distinct from and opposite to that of cortical SST interneurons.

**Figure 4.**
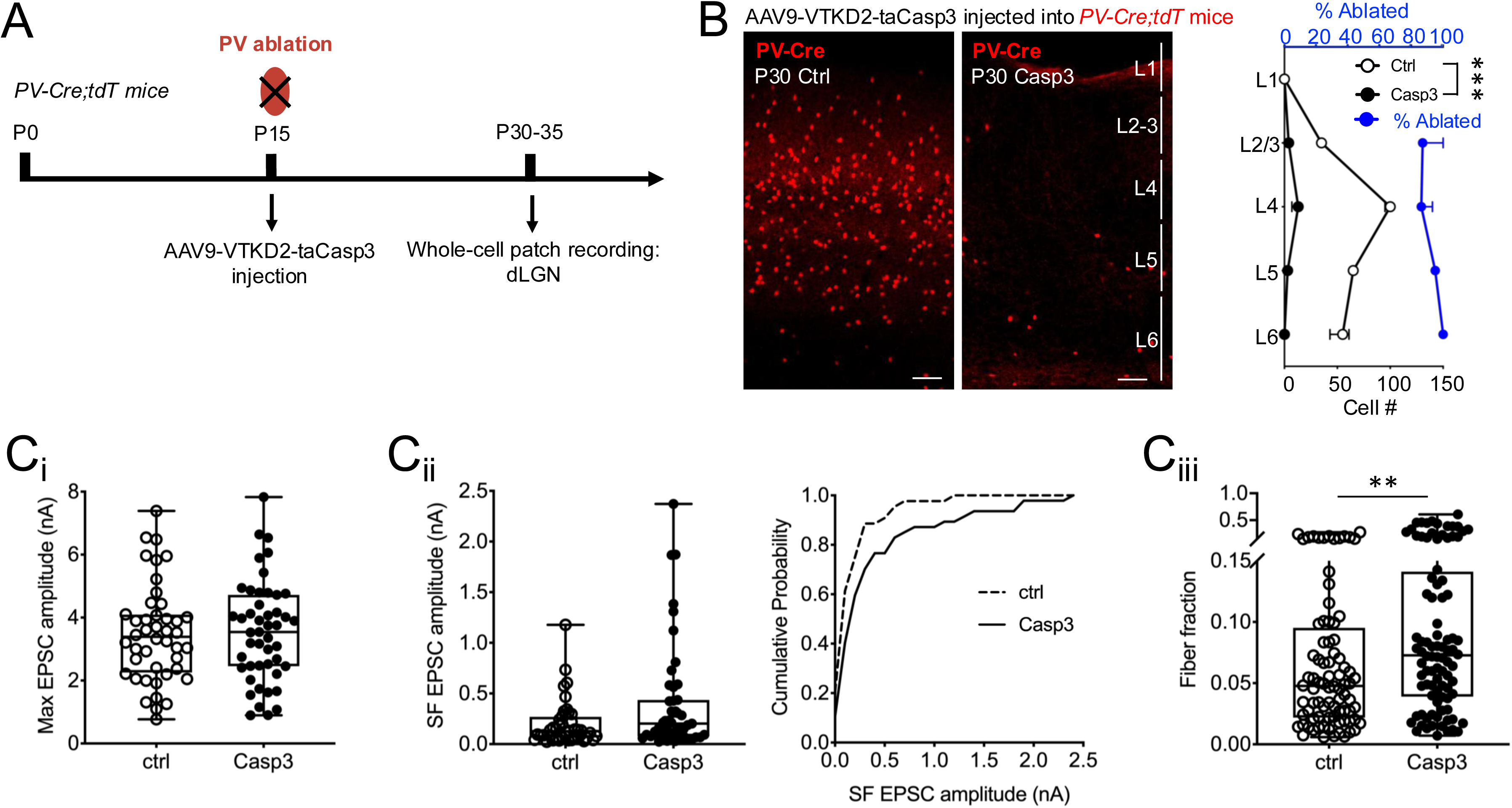
Ablation of cortical PV interneurons during development accelerates retinogeniculate pruning. A. Protocol for ablating cortical PV interneurons. B. Ablation of PV interneurons across different layers in V1. *Left*: representative images of PV-tdT^+^ interneurons in the V1 from ctrl or Casp3-treated mice at P30. *Right*: Distribution of PV-tdT^+^ interneurons across V1 (black axis) and the proportion of PV interneurons ablated by Casp3 (blue axis) (n = 3 mice). *** *P* < 0.001, two-way ANOVA. Scale bar: 100 μm. C. Measurement of retinogeniculate responses after ablation of cortical PV interneurons. C_i_: Peak amplitude of the Max AMPAR EPSCs, *P* = 0.69, Mann-Whitney test; C_ii_: Peak amplitude (*left*) and cumulative probability distribution (*right*) of SF EPSCs. *Left*: *P* = 0.057, Mann-Whitney test; *Right*: *P* = 0.173, Kolmogorov-Smirnov test; C_iii_: Distribution of FF ratio, ** *P* < 0.01, Mann-Whitney test.

### Experience-dependent refinement of retinogeniculate synapses is regulated by cortical SST and PV interneurons

Visual deprivation disrupts both the maturation of cortical SST interneuron-mediated circuits and connectivity at the retinogeniculate synapse. ^1^ This raises the question of whether cortical SST interneurons regulate experience-dependent retinogeniculate plasticity. Therefore, we selectively manipulated cortical SST interneuron activity during LDR. We expressed a Cre-dependent excitatory DREADD (hM3Dq) in cortical SST interneurons at P15 by viral injection, and administered CNO through drinking water between P20 and P30 while the mice were subjected to LDR (see Methods). Control mice underwent the same procedure but received vehicle viral injections instead and were divided into cohorts reared under either normal light/dark cycles or LDR (Fig. 5A). Recordings from these mice show that the amplitude of both maximum and SF EPSCs, as well as the median of FF, are greater in DREADD-manipulated LDR mice compared to control LDR mice. These results show that increased SST interneuron activity prevents both the weakening of single fiber strength and the increase in the number of retinal inputs caused by LDR. In fact, retinogeniculate synapse refinement in LDR and DREADD-manipulated mice is not significantly different from that in normally reared control mice, suggesting that DREADD activation of cortical SST interneurons largely prevents the disrupted retinogeniculate connectivity typically observed in LDR mice (Fig. 5B, Table S3). Building on previous results, our findings demonstrate that LDR impacts SST interneuron circuit maturation, whereas activation of SST interneurons counteracts the impact of LDR on retinogeniculate synapse refinement. These results suggest that developing cortical SST interneurons are functionally modulated by sensory input and, in turn, directly regulate thalamic synapse refinement based on experience.

**Figure 5.**
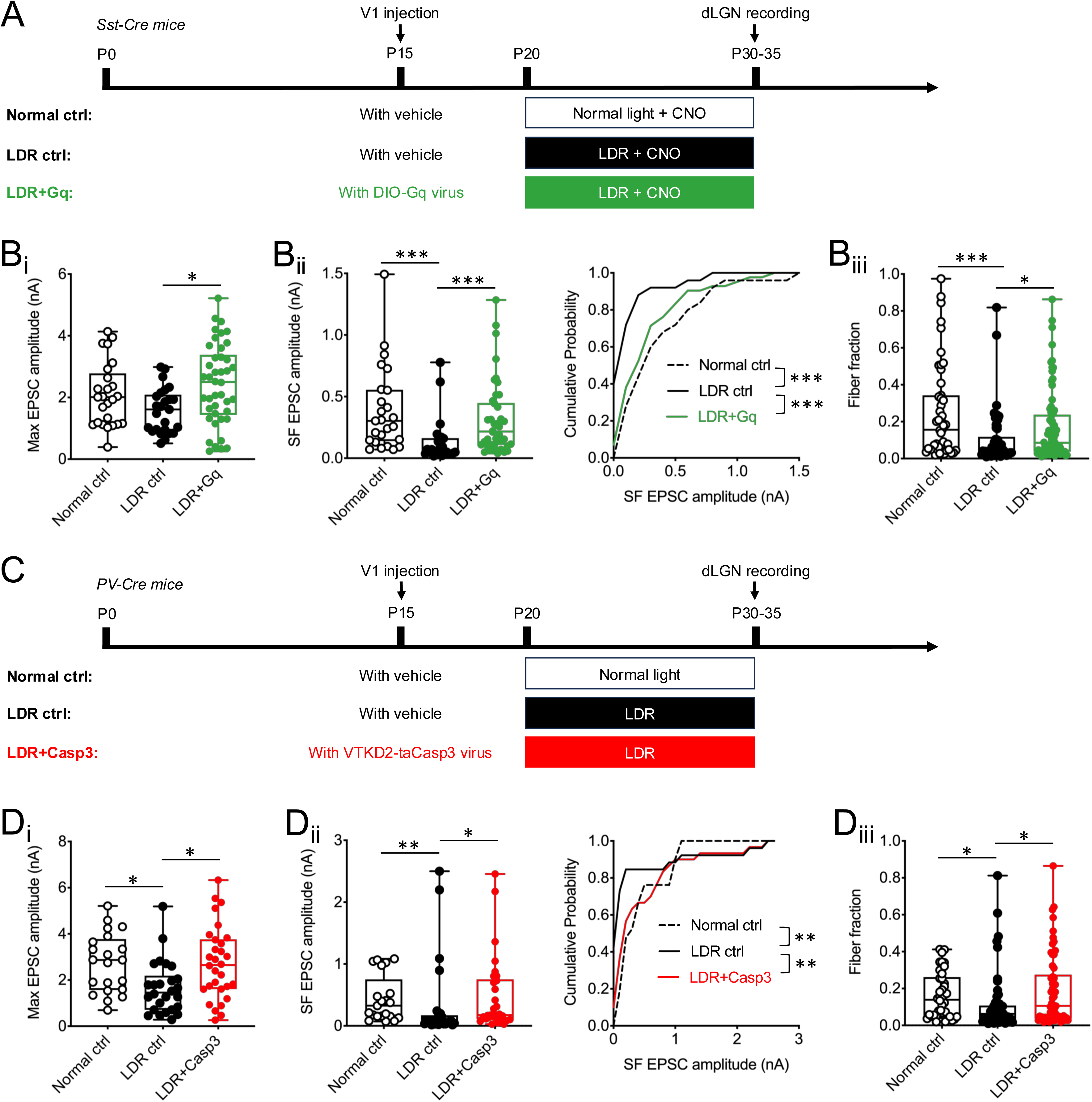
Visual deprivation-triggered retinogeniculate remodeling is prevented by activation of SST or ablation of PV interneurons. A. Time course for activation of SST interneurons and visual deprivation. B. Activation of SST interneurons during P20-30 prevented the alteration in retino-thalamic connection caused by LDR. B_i_: Peak amplitude of the Max AMPAR EPSCs, * *P* < 0.05, Kruskal-Wallis test. B_ii_: Peak amplitude (left) and cumulative probability distribution (right) of SF EPSCs, *** *P* < 0.001, Kruskal-Wallis test and Kolmogorov-Smirnov test, respectively; B_iii_: Distribution of FF ratio, * *P* < 0.05, *** *P* < 0.001, Kruskal-Wallis test. C. Time course for ablation of PV interneurons and visual deprivation. D. Ablation of PV interneurons during P20-30 occludes the remodeling of thalamic refinement triggered by LDR. D_i_: Peak amplitude of the Max AMPAR EPSCs, * *P* < 0.05, Kruskal-Wallis test; D_ii_: Peak amplitude (left) and cumulative probability distribution (right) of SF EPSCs, * *P* < 0.05, ** *P* < 0.01, Kruskal-Wallis test and Kolmogorov-Smirnov test, respectively; D_iii_: Distribution of FF ratio, * *P* < 0.05, Kruskal-Wallis test.

Our finding that PV ablation enhances pruning raises the question of whether PV interneurons also directly participate in sensory-dependent synaptic refinement. To explore this, we ablated PV interneurons in mice subjected to LDR and measured retinogeniculate connectivity between P30-35 (Fig. 5C). Remarkably, similar to the effects observed when SST interneurons are activated, both SF strength and FF are significantly increased in PV-ablation mice compared to LDR controls, reaching levels comparable to those in normally reared control mice (Fig. 5D, Table S3). These results suggest that cortical SST and PV interneurons provide complementary, bidirectional control over retinogeniculate remodeling, regulating synaptic refinement in response to visual experience.

### SST interneurons modulate the maturation of PV interneuron firing properties

Up to this point, we have examined the overall effects of SST and PV interneurons by manipulating them separately, demonstrating their opposing effects on the refinement of retinogeniculate synapses. Yet, whether there is interaction between these two types of interneurons is not clear. Given that SST interneurons mature earlier and form increasingly stronger synapses onto PV interneurons during development (Fig. 1C), this interaction may play a role in balancing their opposing influences on thalamic synapse refinement. Our results showed that the developmental strengthening of SST→PV synapses does not appear to be influenced by sensory experience (Fig. 2B). To investigate further, we tested whether SST interneurons impact other aspects of PV interneuron maturation in a sensory experience-dependent manner. Given the late onset of PV expression, we first examined whether PV expression correlates with the electrophysiological maturation of PV interneurons. By recording from PV-tdT^+^ neurons in L6 of *Pvalb-tdT* mice, we observe heterogeneity in their firing patterns in response to current injections between P15 and P30 (see Methods). We identify three distinct firing patterns: single-spiking, slow-spiking (with a maximum firing rate below 40 spikes/s, Fig. S5A), and fast-spiking (with a maximum firing rate above 50 spikes/s, Fig. S5B). At P15, only 16% of tdT^+^ cells are fast-spiking, while the majority are single- or slow-spiking (Fig. 6A). The fraction of fast-spiking tdT^+^ cells increases to 65% at P20, and reaches 100% at P30 (Fig. 6A), accompanied by a developmental reduction of the action potential half-width, rise and decay times (Fig. S5C-E). Therefore, PV interneurons gradually acquire their mature firing properties over an extended developmental period before P30. In contrast, SST interneurons assume their mature spiking properties much earlier, before P22-24. ^37^

**Figure 6.**
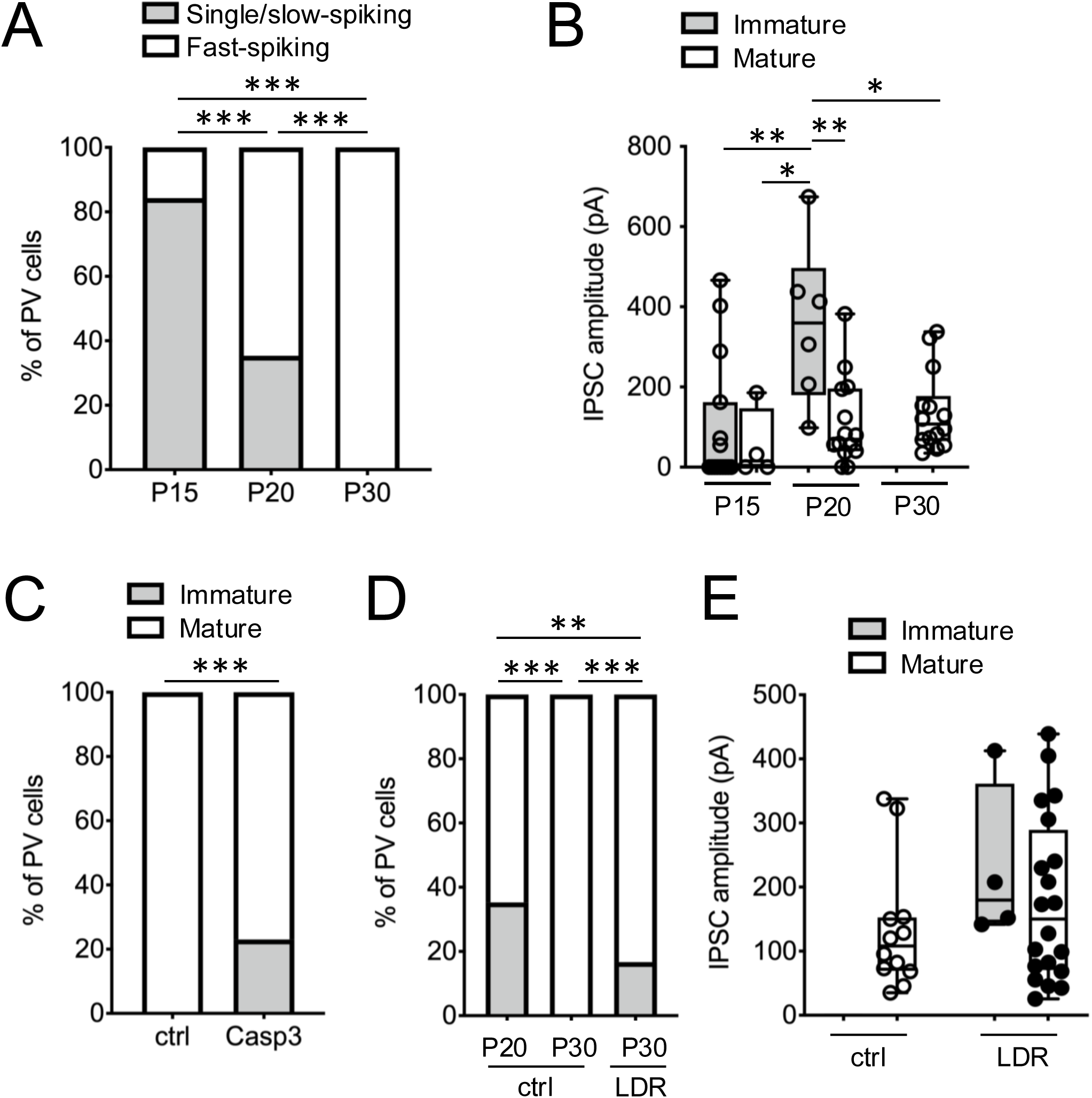
SST interneurons modulate the maturation of PV interneuron firing properties. A. Fractional distribution of single to slow-spiking vs fast-spiking PV interneurons during development. *** *P* < 0.001, Chi-square test. B. Amplitude of SST→PV IPSCs in immature (single/slow-, gray bars) vs mature (fast-, white bars) PV interneurons during development. * *P* < 0.05, ** *P* < 0.01, two-way ANOVA, Sidak’s multiple comparisons test. C. Fractional distribution of immature vs mature PV interneurons after ablation of SST interneurons. *** *P* < 0.001, Chi-square test. D. Fractional distribution of immature vs mature PV interneurons after LDR. ** *P* < 0.01, *** *P* < 0.001, Fisher’s exact test. E. Median amplitude of SST→PV IPSCs sorted by maturation state of PV cells after LDR.

To explore how the maturation of PV interneuron electrophysiological properties is linked to developmental changes in the SST→PV synapse, we grouped the SST→PV synaptic responses into two categories according to PV firing properties: single to slow-spiking (immature) versus fast-spiking (mature). We find that the median amplitude of SST-evoked IPSCs onto single to slow-spiking PV cells significantly increases between P15 and P20 (Fig. 6B, Table S2), consistent with stronger SST synaptic charge transfer onto immature PV interneurons at P20 (Fig. S5F). In contrast, the strength of SST input onto fast-spiking PV interneurons is weaker and does not change significantly between P15 and P30 (Fig. 6B, Table S2). These results are consistent with a transient strengthening of SST→immature PV synapses during development, followed by a net reduction in the average strength of all SST→PV inputs (both immature and mature PV interneurons) as PV interneurons become fast-spiking. The strong inhibition of immature PV interneurons might serve to promote the maturation of PV interneurons, or to prevent the ‘brake’ on pruning from occurring prematurely.

To investigate how SST interneurons influence the maturation of PV firing properties, we repeated the ablation of SST interneurons as before. While the number of PV interneurons remains unchanged after SST interneuron ablation (Fig. S3B), 23.1 % of these PV interneurons exhibit single to slow-spiking patterns at P30 (Fig. 6C). This finding underscores the significant role of SST interneurons in the final maturation of PV interneuron electrophysiological properties. Intriguingly, similar fraction (∼17 %) of PV cells exhibit single to slow-spiking pattern at P30 after LDR (Fig. 6D, E). These results suggest that SST interneurons promote the electrophysiological maturation of PV interneurons in response to sensory input. By doing so, they help activate PV interneurons, which then acts as a counterbalancing force alongside SST interneurons to fine-tune the retinogeniculate synapse pruning.

## Discussion

There is growing evidence supporting the idea that experience-dependent plasticity engages bidirectional circuits between the thalamus and primary visual cortex (V1). ^5,38,39^ Circuit refinement within these two visual regions does not occur independently and sequentially, but rather entails interactions between feedforward and feedback projections. ^40–46^ In the present study, we identify key components of cortical feedback circuits responsible for vision-dependent refinement of feedforward retinogeniculate connectivity during late development. Here we offer a working model based on our results which can serve as a framework upon which future studies can refine our understanding of cortico-thalamic interactions during development.

### A reciprocal antagonistic mechanism of cortical inhibition contributes to top-down regulation of retinogeniculate refinement

The most parsimonious interpretation of our results is that SST interneurons in V1 contribute to driving the strengthening and pruning of the retinogeniculate synapse as both cortical and thalamic circuits remodel to incorporate visual experience into their connectivity. In contrast, the continued maturation of cortical PV circuits plays a role in limiting retinogeniculate pruning, by slowing down or “braking” the consolidation of thalamic circuits during the sensitive period so that experience-dependent changes can still be made. ^47–50^ The balance between the two opposing cortical circuits shift over the course of the sensitive period such that the influence of PV circuits increase with age and retinogeniculate connectivity stabilizes by the end of the period.

We propose that this shift in balance is coordinated by SST interneuron-mediated maturation of PV neuron spiking. Supporting this model, our findings reveal that SST interneurons develop much earlier than PV interneurons. As they mature, they shape the development of the infragranular circuits within V1 and promote retinogeniculate refinement by increasing inhibition influence on the activity of L6 PN neurons. Our previous study demonstrated that distinct manipulations of L6 corticothalamic neuron activity at P20 can differentially alter the strength or number of convergent retinal inputs onto TC neurons. ^5^

This model is also consistent with our findings that with SST interneuron ablation (and presumably also LDR) there is reduced inhibition onto L6 PNs, shifting the balance of forward circuit maturation and PV circuit “braking” towards the latter, leading to greater number of convergent retinal inputs that are weaker. Conversely, ablation of PV interneurons leads to a shift of the reciprocal balance towards accelerated refinement of the retinogeniculate synapse (Fig. 7). Taken together, our results support a critical role for SST interneurons in initiating the forward driving of thalamic refinement.

**Figure 7.**
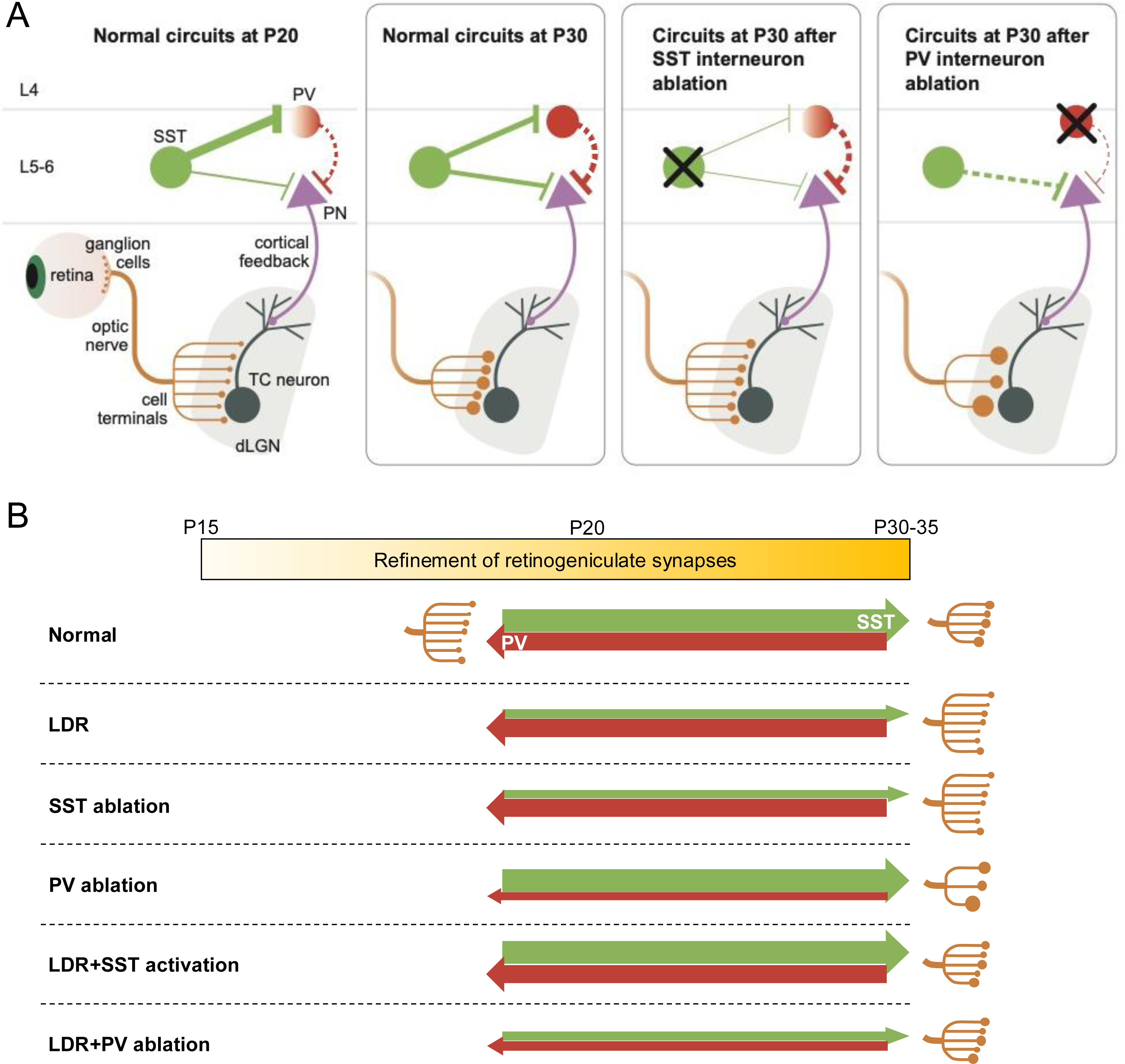
Reciprocal regulation of retinogeniculate refinement by cortical SST and PV interneurons. A. Summary of effects of ablating cortical SST vs. PV interneurons on the refinement of retinogeniculate synapses in the dLGN. The thickness of the green/red lines in the V1 indicates relative strength of inhibition from SST/PV interneurons. Dashed lines represent speculated changes in the strength of connectivity. In normal development from P20 to P30, when retinal inputs undergo experience-dependent strengthening and pruning, cortical SST interneurons mediate the maturation of infragranular inhibitory circuits. When SST interneurons are ablated between P20 and 30, PV interneurons take over the role as being the main inhibitor of PNs, the retinogeniculate synapses are much less refined in the dLGN. On the contrary, loss of PV interneurons between P20-30 leads to acceleration in the pruning of retinal inputs compared to normal circuits in the thalamus. B. Schematic for the opposing forces between cortical SST driving forward development (green arrows) and PV interneurons braking refinement (red arrows). The thickness of the arrow bars indicates relative strength from driving vs. braking refinement. During normal development, SST and PV interneurons form a balance in the driving vs. braking system to determine the timing of strengthening and pruning of retinal inputs over age. After LDR or ablation of SST interneurons, the balance of forward vs. braking refinement shifts, leading to less refined thalamic circuits. On the contrary, with PV interneuron ablation, the brakes are removed, allowing thalamic circuits to develop more rapidly. Enhanced SST interneuron activity overrides LDR effects, while PV ablation during LDR establishes a new balance between the opposing drives of SST and PV interneuron.

### Interactions between SST and PV interneuron circuits over development

Our results are consistent with a growing body of evidence demonstrating that the maturation of cerebral cortex is regulated by the interplay of SST and PV interneurons. Work in both the somatosensory ^15,18,22,51,52^ and the visual system ^16,17,19^ indicates that the transition from synchronous to decorrelated activity in the cerebral cortex involves a relay involving an early role of SST shifting to PV activity. ^13,20–23^ This results in a sparsification of cortical activity, which we here suggest sequentially alters top-down signaling to impact retinogeniculate development. These results thus provide further insight into how the relationship between SST and PV circuits impacts thalamic development. The fact that ablation of PV interneurons displays an opposite outcome in synaptic pruning to that when SST interneurons are ablated rules out a simple feedforward model where PV interneurons act solely as a downstream mediator of SST interneurons. Instead, the functional output of the circuitry between SST and PV interneurons likely involve network interactions in the cortex, as described above. In particular, the results from ablation of PV interneurons in LDR mice are illuminating. If the influence on retinogeniculate refinement were dependent on the absolute level of activity of SST vs. PV interneurons, it would be hard to explain how reduction of both SST and PV interneuron drive leads to normal retinogeniculate connectivity. Instead, we suggest that it is the balance of SST vs. PV interneuron circuit drive that is important (Fig. 7B). Yet SST and PV interneuron circuits are not fully independent, as we find that LDR or ablation of SST interneurons at around P20 prevents the final maturation of PV interneuron excitability. Therefore, full cortical influence on retinogeniculate refinement depends on both independent SST and PV interneuron circuits as well as complex interactions between these circuits that have been recently demonstrated. ^22^

### Experience-dependent plasticity of the corticothalamic circuit

Our finding that the sensitivity to visual experience of SST circuits mirrors that of the retinogeniculate synapse highlights a distinct relationship between these specific cortical and thalamic circuits. These findings are different from previous reports of experience-dependent refinement of other visual cortical circuits. Complete deprivation of visual experience from birth (referred to as chronic dark rearing, CDR) is known to prevent the maturation of visual acuity and development of PV interneuron circuits. ^53–56^ Studies have demonstrated that prolonged deprivation decreases GAD65- and GAD67-immunopositive perisomatic puncta on pyramidal neurons and PV expression. ^57–60^ In contrast, neither the density of SST^+^ interneurons nor the expression of SST is changed in the cortex by CDR. ^54,57^ Dark rearing from P20 (LDR) is a very different manipulation from CDR as much of the inhibitory circuits have already developed by this time. Moreover, we find that SST➝PN synaptic transmission is not sensitive to CDR (Fig. S2E). Notably, our previous studies showed that CDR did not elicit the same retinogeniculate plasticity as LDR –– instead retinogeniculate refinement appeared normal. ^1,2^ Taken together, our results demonstrate that different cortical inhibitory circuits exhibit sensitivity to visual experience during distinct windows of development.

While our working model of synaptic refinement in the geniculate is consistent with cortical SST interneurons accelerating and PV interneurons attenuating synaptic refinement, intracortical circuits are complex. Although our previous evidence demonstrates that retinogeniculate remodeling can be modulated by alterations in corticothalamic feedback, ^5^ it remains unclear how this pathway is controlled by interneuron dynamics. Notably, these events occur subsequent to when cortical activity changes from synchronous to decorrelated, events thought to be dependent upon SST to PV maturation. ^13,15–23^ Taken together, this may indicate a higher-order role for how cortical networks impact top-down signaling through SST and PV interneuron maturation.

## Acknowledgments

We thank Drs. April Levine, Brielle Ferguson, John Assad, and all other members of the Chen lab for helpful discussions about the project and manuscript. We thank Jianlin Wang, Israel Robinson and Iris Wu for blinding the experimenter, and Shuhan Huang from Dr. Gord Fishell’s lab for providing the *AAV9-VTKD2-taCasp3* virus. Support was provided by the NIH RO1EY013613 and Tan-Yang Center for Autism Research Grant to C.C., William Randolph Hearst Fund 520.45318.7900.600377.730002.0000.65852 to Q.J., and NIH 5R01NS081297 to G.F.. We thank the IDDRC Cellular Imaging Core, funded in part by S10OD016453 for access to their shared confocal microscopes, as well as P50HD105351 for the Cellular Imaging and Administrative cores.

## Author contributions

Q.J., G.F. and C.C. designed the experiments and wrote the paper. Q.J. conducted the whole-cell patch recordings from both V1 and dLGN neurons. S.W. performed RNAscope experiment.

## Materials and Methods

### Animals

All animal procedures were in compliance with the NIH Guide for the Care and Use of Laboratory Animals and approved by the Institutional Animal and Care and Use Committee (IACUC) at Boston Children’s Hospital. To label and drive SST interneurons in the V1, *Sst-IRES-Cre* (*Sst-Cre*) transgenic mice (JAX 013044) were crossed with fluorescently-tagged Cre-dependent tdTomato expressing mice (*Ai14*, JAX 007908), ^61^ or ChR2-EYFP expressing mice (*Ai32*, JAX 012569). ^62^ We refer to the resulting crosses as “*Sst;tdT*” and “*Sst;ChR2*”, respectively. *Sst;ChR2* mice were also crossed with *Pvalbumin-tdTomato* (*Pvalb-tdT*) transgenic mice (JAX 027395) to yield *Pvalb;Sst;ChR2* progenies for examining SST inhibitory transmission onto PV interneurons in V1. *Pvalb-Cre* mice were obtained from Dr. Clifford Woolf’s lab to ablate PV interneurons specifically (JAX 017320). ^63^ They were crossed with *Ai14* mice to yield *Pvalb-Cre;tdT* progenies to detect the developmental distribution of PV-Cre^+^ interneurons in the cortex. Mice aged P10-P60 of either sex were used. For chronic CNO treatment, mice had ad libitum access to CNO-treated water (0.25 mg/ml) instead of regular drinking water.

### Visual deprivation

For late dark rearing, mice were subjected to dark rearing during P20-(∼30). At desired ages, they were sacrificed in the dark for slice preparation. For chronic dark rearing, the pups were placed in dark boxes right after birth together with their mother and sacrificed in the dark without exposure to normal light.

### Tissue preparation and immunohistochemistry

Mice were anesthetized with 50 mg/kg pentobarbital and transcardially perfused with 0.1 M phosphate buffered saline (PBS) immediately followed by 4% w/v paraformaldehyde (PFA) in PBS. Brains were post-fixed overnight in 4% PFA at 4°C and rinsed in PBS. Brain slices containing V1 were coronally sectioned through Leica VT1000 vibratome with thickness of 60 μm.

For immunostaining, brain slices containing V1 were blocked in PBS containing 5% normal goat serum (NGS) and 0.1% Triton X-100 at room temperature for 1 hr. Then primary antibodies were applied in PBS containing 0.1% Triton and 2% NGS: chicken anti-GFP (1:1000; ab13970, Abcam), and/or rabbit anti-RFP (1:1000; 600-401-379, Rockland), and/or rabbit anti-PV (1:1000; PV27, SWant) at 4°C for overnight. After rinsing with 0.1% Triton/PBS in the next day, the slices were incubated with secondary antibodies at room temperature for 2 hours: goat anti-chicken antibody conjugated to Alexa Fluor 488 (1:1000; A11039, Invitrogen), and/or goat anti-rabbit antibody conjugated to Alexa Fluor 488 (1:1000; ab150077, Abcam), or 555 (1:1000; A32732, Invitrogen). Slices were then incubated with DAPI for nuclear detection, mounted and cover-slipped with Vectashield (VectorLabs H-1000). For the quantification of V1 corticothalamic neurons, a combination of primary antibodies: rabbit anti-GABA (1:1000; A2052, Sigma), mouse anti-Tle4 (1:100; sc-365406, Santa Cruz), and chicken anti-NeuN (1:1000; ABN91, Millipore Sigma); and secondary antibodies: goat anti-rabbit Alexa 647 (1:1000; A21245, Invitrogen), goat anti-mouse Alexa 555 (1:1000; A32727, Invitrogen), and goat anti-chicken Alexa 488 (1:1000; A11039, Invitrogen) were used for the staining of slices containing V1.

### Single Molecule Fluorescent *In Situ* Hybridization Histochemistry

For single molecule fluorescent in situ hybridization (smFISH) combined with immunohistochemistry, mice were perfused and brains were fixed overnight in 4% PFA in 1 x PBS followed by cryoprotection in 30% sucrose in 1 x PBS. Then, 16-20 μm thick brain sections were obtained using a Leica cryostat. The sectioned brain slices were directly mounted on glass slides (Fisherbrand Superfrost Plus) and preserved in -80°C freezer.

For RNAscope experiments, samples were processed according to the ACDBio Multiplex Flourescent v2 Kit protocol (ACDBio #323100) for fixed frozen tissue. Briefly, tissue was pre-treated with a series of dehydration, H_2_O_2_, antigen retrieval and protease III steps before incubation with the probe for 2 hours at 40°C. Note here protease III incubation was performed at room temperature to better preserve the protein for immunostaining. The probes used for labeling included 1) *RNAscope Probe-Mm-Pvalb* (Cat#421931-C3, ACDBio); and 2) *RNAscope Probe-Mm-Gad1* (Cat#400951, ACDBio). Three amplification steps were carried out prior to developing the signal with Opal™ or TSA® Dyes (Akoya Biosciences). Immuostaining following RNAscope experiment was performed according to Technical Note 323100-TNS from ACDBio. Samples were counterstained with DAPI and mounted using Prolong Gold antifade mounting medium (Molecular Probes #P369300).

### Imaging and image analysis

To measure the expression of SST-tdT^+^, PV-tdT^+^, PV^+^, or PV-Cre^+^ neurons over development or over time after injection of viruses, images of the V1 (one coronal full view containing both monocular and binocular regions from each animal) from at least three mice were acquired with Zeiss LSM 700 (Zeiss, Olympus) using a 10x objective. Scans were performed to obtain 9 to 11 optical Z sections of 6 μm each. Quantification was performed manually using ImageJ. The images were stacked over the whole slice with thickness of ∼60 μm. The number of tdT^+^ or PV^+^ neurons were then counted manually from each layer in the V1, including L1, L2/3, L4, L5 and L6. For the identification and measurement of single molecule signals of *Pval* and *Gad1* following RNAscope experiment, or the assessment of overlap among molecules of Tle4, NeuN and GABA, V1 images were acquired through Zeiss LSM 700 using 40x and 63x objectives with built-in functions of Z-stack. Consecutive images (0.9 μm thick each) were collected for analysis. Maximum projection and cell counting were conducted through ImageJ. In the *RNAscope* images, a cluster of single molecules was identified as belonging to one cell when they co-localized with DAPI and formed a clear cell morphology.

### Viral injections

Mice were anesthetized via 2% of isoflurane (Baxter, Illinois) and fixed onto the stereotactic platform. Viruses expressing Caspase3 (Casp3) were microinjected using NanojectIII (Drummond, Pennsylvania) at 1 nL/second to three sites of the right hemisphere of V1 to cover both monocular and binocular zones. The following coordinates were used: P15: from Lambda AP+0.7, ML+2.4, DV-0.55; AP+1.4, ML+2.7, DV-0.55; AP+0.7, ML+2.9, DV-0.55; P30: from Lambda AP+0.8, ML+2.35, DV-0.55; AP+1.5, ML+2.35, DV-0.55; AP+0.8, ML+2.95, DV-0.55 (in mm). Virus *AAV9-VTKD2-taCasp3-TEVp* (shared by Gord Fishell’s lab) was used for ablation of SST or PV interneurons, and *AAV2/9-hSyn-DIO-hM3D(Gq)-mCherry* (Addgene #44361) for activation of SST interneurons. The mice were perfused for immunostaining or sacrificed for patch recording at required ages. Control (ctrl) mice and those injected with Casp3-expressing virus were housed together and randomly assigned to the experimenter for patch recording as a blind of condition.

### Electrophysiology

Brain slices for *in vitro* recordings were prepared as previously described. ^64,65^ Briefly, mice were anesthetized using isoflurane and decapitated into oxygenated (95% O_2_; 5% CO_2_) ice-cold cutting solution (in mM): 130 K-gluconate, 15 KCl, 0.05 EGTA, 20 HEPES, and 25 glucose (pH 7.4 adjusted with KOH, 310-315 mOsm). ^66^ The brain was then removed quickly and immersed in the ice-cold cutting solution for 60 seconds. For V1 recording, coronal slices containing V1 were sectioned and collected. For dLGN recording, parasagittal sectioning was conducted to obtain slices maintaining continuity of optic tracts (OT) as previously described. ^67^ The brain was cut with a steel razor blade, then sectioned into 250 μm-thick slices in the oxygenated ice-cold cutting solution using a sapphire blade (Delaware Diamond Knives, Wilmington, DE) on a vibratome (VT1200S; Leica, Deerfield, IL). The slices collected were allowed to recover at 30°C for 15-20 minutes in oxygenated saline solution (in mM): 125 NaCl, 26 NaHCO_3_, 1.25 NaH_2_PO_4_, 2.5 KCl, 1.0 MgCl_2_, 2.0 CaCl_2_, and 25 glucose (pH 7.4, 310-315 mOsm).

For cortical recordings, PV interneurons or PNs in V1 L6 were visualized through a monitor with projection from the camera of a DIC-equipped microscope (Prime BSI, Teledyne Photometrics). While PV interneurons were genetically labeled in *Pvalb-tdT* mice, PNs were identified based on their distinct pyramid-like shape and relatively larger size compared to interneurons. L6 was identified as no more than 200 μm away from the white matter. Glass pipettes (Drummond Scientific) were pulled on Sutter P-97 Flaming/Brown micropipette puller (Sutter Instruments) and filled with internal solution containing (in mM): 150 K-gluconate, 8 KCl, 10 EGTA, 10 HEPES (pH7.3, 290-300 mOsm) to optimize the pipette resistance to be 3.5-4.0 MOhm. Patch recordings were performed using a MultiClamp 700B (Axon Instruments, Foster City, CA) and an ITC-18 interface (Instrutech) with sampling rate of 2 kHz and filtering frequency of 5 kHz. Inhibitory post-synaptic currents (IPSCs) were obtained by holding the membrane potential at 0 or 30 mV and applying a single pulse (0.2 ms) of full-field illumination of blue light (470 nm) through the 60x objective (Olympus LUMplanFL N 60x/1.00W), thus to confine the illumination to an area with a radius of ∼220 μm within the infragranular layers without surpassing L4. The blue light was supplied by a CoolLED pE unit, lasting for 0.2 msec at highest power (100%, 83 mW/mm^2^) to obtain maximal current. Intertrial intervals were kept at 1 min. Access resistance was monitored throughout the experiment and evaluated in offline analysis. Experiments with access resistance changing over 20% were removed from analysis. Spiking of SST interneurons and intrinsic cellular properties of PNs and PV interneurons were measured in current clamp mode. I-V curves were obtained by recording firing rates when using current injection from 0 to 600 pA in steps of 50 pA. The activation of SST interneurons was verified by continuous blue light illumination for 1 s, or single pulse (0.2 ms) photo-stimulation in current clamp mode.

To measure retinogeniculate refinement, TC neurons located in the ventral posterior region of the dLGN were recorded as previously described. ^1,68^ Glass pipettes were filled with internal solution containing (in mM): 35 CsF, 100 CsCl, 10 EGTA, 10 HEPES, and L-type calcium channel antagonist 0.1 methoxyverapamil (pH7.3, 290-300 mOsm) to optimize the pipette resistance to be 1.5-2.0 MOhm. Both AMPAR and NMDAR currents were obtained by holding the membrane potential of recorded cells at -70 and +40 mV, respectively. To isolate excitatory synaptic currents, cells were recorded at room temperature in oxygenated saline solution containing 20 μM of bicuculline (GABA_A_R antagonist), 2 μM of CGP55845 (GABA_B_R blocker), 10 μM of DPCPX (antagonist of A1 adenosine receptors), and 50 μM of LY341495 (blocker of presynaptic mGluRs). ^69–73^ To obtain maximal electrical stimulated excitatory post-synaptic currents (EPSCs), a pair of electrodes were filled with saline solution, and lowered onto the slices. One of the electrodes was inserted into the optic tract to electrically stimulate the retinogeniculate inputs. The other electrode was immersed in the bath just above the brain slice surface, serving as the ground. Electrical stimuli were supplied by a stimulus isolator (WPI A365) delivering a 0.2 msec pulse between 0-99 mA. Maximal currents were defined as the largest response that does not increase with higher stimulating intensity (up to 99 mA). Single fiber strength was defined as the first consistent response observed after an increase in stimulation intensity by 0.25 µA. Fiber fraction (FF) was calculated as single fiber strength over maximal current, as an estimate of the contribution of a single input to the total retinal drive, as well as the number of inputs onto one TC neuron. AMPAR currents were included in the analysis of maximal currents and single fiber input, while both AMPAR and NMDAR currents were used for the calculation of FF.

### Data analysis and statistics

Electrophysiological data acquisition and offline analysis were performed using custom software in IgorPro (Wave-Metrics, Portland, OR). EPSC and IPSC amplitudes were obtained from average traces of 3-5 trials. Data calculation and statistical analysis were conducted using Prism (GraphPad Software, Inc.) and MATLAB_R2019b (Mathworks). All data sets were evaluated for normality using the Kolmogorov-Smirnov test. For nonparametric distributions, the Mann-Whitney or Kruskal-Wallis test were used for comparisons between two or among multiple groups. For normally distributed data sets, the Student’s t-test or one-way ANOVA were used. For comparison of time series repeated measurements, two-way ANOVA test was used. All data were presented as medians (interquartiles (IQR)). The box and whisker graphs indicate the median (line within box), 25-75% quartile range (box), and minimum and maximum range (whiskers). For all figures, * *P* < 0.05; ** *P* < 0.01; *** *P* < 0.001.

