## Supplementary material for "Reciprocal interaction between cortical SST and PV interneurons in top-down regulation of retinothalamic refinement": Fig. S1

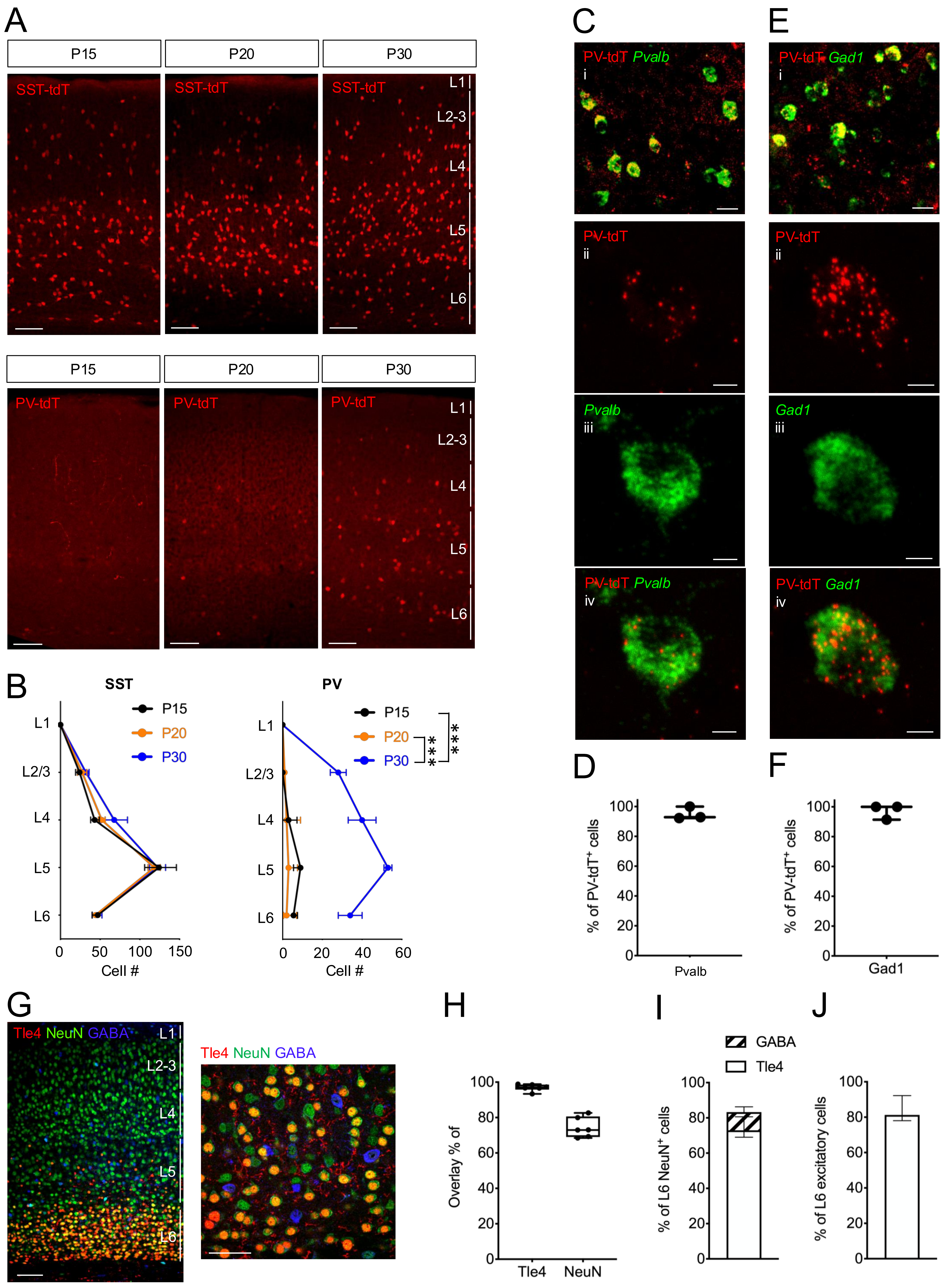

A
P15
P20
P30
SST-tdT
L1
L2-3
L4
L5
L6
PV-tdT
B
SST
PV
P15
P20
P30
L1
L2/3
L4
L5
L6
0
50
100
150
20
40
60
Cell #
***
***
C
PV-tdT Pvalb
i
ii
iii
iv
E
PV-tdT Gad1
D
F
% of PV-tdT+ cells
Pvalb
Gad1
G
Tle4 NeuN GABA
H
Overlay % of
Tle4
NeuN
I
GABA
Tle4
% of L6 NeuN+ cells
J
% of L6 excitatory cells

Figure S1. SST interneurons develop earlier than PV interneurons in the V1, related to Figure 1.

A. Representative images of SST (*top*) and PV (*bottom*) interneurons over development. Scale bar: 100 μm.

B. Developmental expression of SST (*left*) and PV interneurons (*right*) across different layers of V1. SST: P15: n = 3; P20: n = 3; P30: n = 3; PV: P15: n = 6; P20: n = 9; P30: n = 3. *Left*: The number of SST interneurons does not alter between P15-30. $P = 0.39$, two-way ANOVA test. *Right*: There is a significant increase in the number of PV-tdT$^+$ interneurons across different layers from P20 to P30. *** $P < 0.001$, two-way ANOVA test.

C. *In situ* hybridization staining of PV-tdT$^+$ interneurons with RNA probe of *Pval*. $C_i$: Confocal images of co-labeling between tdT and *Pval*. Scale bar: 20 μm; $C_{ii\text{-}iv}$: co-labeling of one PV-tdT$^+$ neuron with *Pval*. Scale bar: 5 μm.

D. Proportion of PV-tdT$^+$ interneurons that are labelled by *Pval* (n = 3 mice).

E. *In situ* hybridization staining of PV-tdT$^+$ interneurons with RNA probe of *Gad1*. $E_i$: Confocal images of co-labeling between tdT and *Gad1*. Scale bar: 20 μm; $E_{ii\text{-}iv}$: co-labeling of one PV-tdT$^+$ neuron with *Gad1*. Scale bar: 5 μm.

F. Proportion of PV-tdT$^+$ interneurons that are labelled by *Gad1* (n = 3 mice).

G. Labeling of Tle4$^+$ and GABA$^+$ neurons in V1 at P30. *Left*: Confocal image of V1 labeled with antibodies to Tle4, GABA, and NeuN across all layers. Scale bar: 100 μm; *Right*: Magnified image from V1 L6. Scale bar: 50 μm.

H. Percentage of Tle4$^+$ cells that are co-labeled by NeuN , and percentage of neurons (NeuN$^+$) that are Tle4$^+$ in V1 L6.

I. Percentage of corticothalamic (Tle4$^+$) and inhibitory (GABA$^+$) neurons in V1 L6.

J. Percentage of corticothalamic (Tle4$^+$) neurons based on excitatory neurons in V1 L6. The number of excitatory neurons was calculated by subtracting the number of GABA$^+$ from NeuN$^+$ cells. G-J: n = 6 mice.
