## Supplementary material for "Reciprocal interaction between cortical SST and PV interneurons in top-down regulation of retinothalamic refinement": Fig. S2

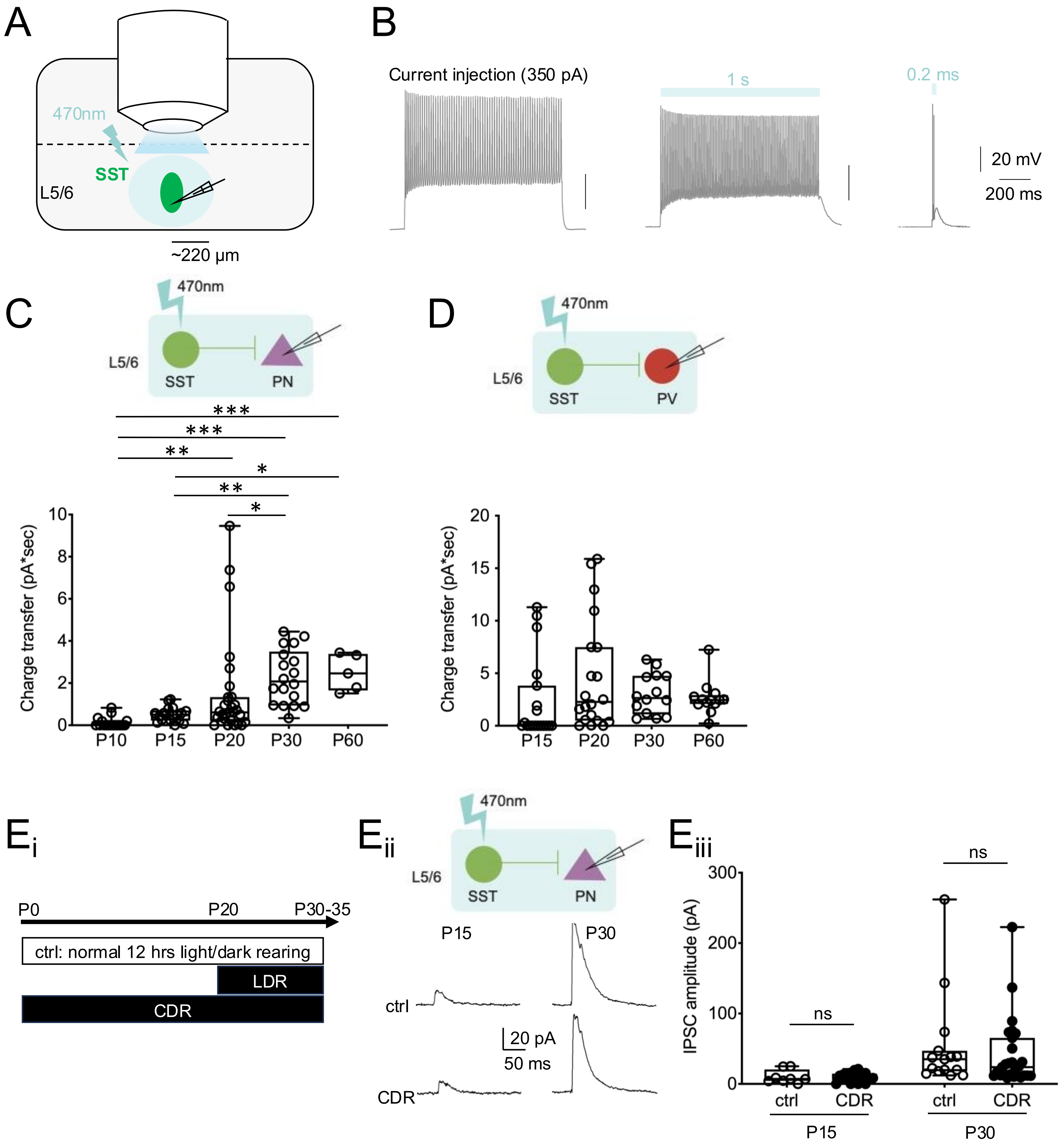


Figure S2. SST interneuron-mediated inhibitory circuits mature before the end of thalamic sensitive period, and are not sensitive to chronic dark rearing (CDR), related to Figures 1 and 2.
A. Schematic of photostimulation of SST interneurons in V1 L5/6 through 60x objective to confine the area within a radius of ~220 μm (See Methods).
B. Example traces of firing from one SST interneuron in response to current injection of 350 pA (*left*, minimum injected current triggering highest firing frequency), continuous blue light stimulation (1 s) (*mid*), and single pulse of stimulation (0.2 ms) (*right*). Blue light illumination effectively activates spiking of ChR2-expressing SST interneurons comparable to that with current injection.
C. Developmental time course for the charge transfer of SST→PN IPSCs.
D. Developmental time course for the charge transfer of SST→PV interneuron IPSCs. C-D: * $P < 0.05$, ** $P < 0.01$, *** $P < 0.001$, Kruskal-Wallis test, Dunn's multiple comparisons test.
E. Developmental change of SST→PN IPSCs under CDR. $E_i$: Schematic of LDR and CDR. $E_{ii}$: Example traces; $E_{iii}$: Peak amplitudes of SST→PN IPSCs at P15 and P30. P15: $P > 0.99$; P30: $P > 0.99$, two-way ANOVA, Sidak's multiple comparisons test.
