## Supplementary material for "Reciprocal interaction between cortical SST and PV interneurons in top-down regulation of retinothalamic refinement": Fig. S3

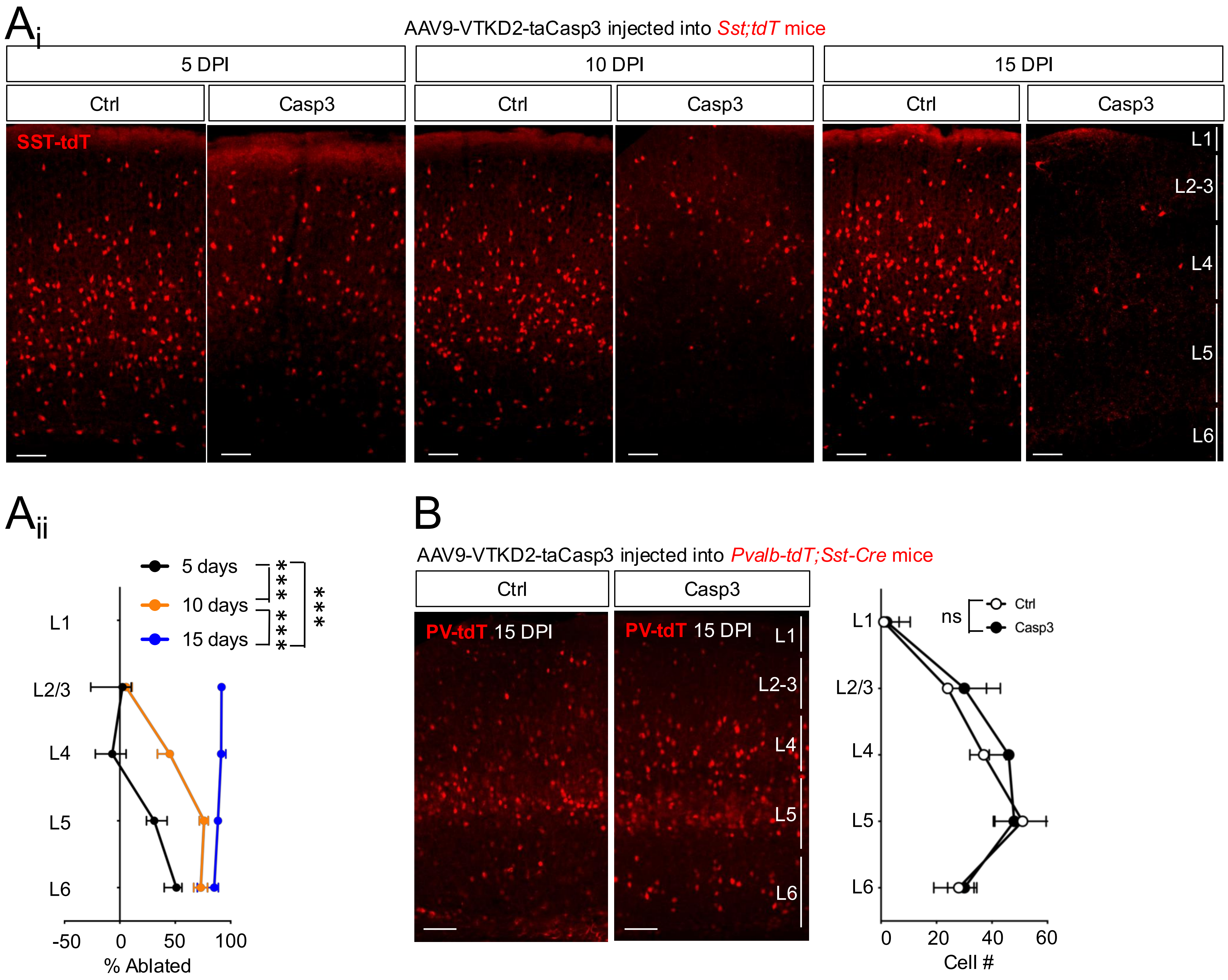


Figure S3. Ablation of SST interneurons in the V1 is detectable by 5 DPI, related to Figure 3.

A. Distribution of SST interneurons with time after expression of Casp3 at 5 (n = 3), 10 (n = 3), and 15 (n = 8) days post-injection (DPI). $A_i$: Representative images of remaining SST interneurons in the V1 after 5, 10, or 15 DPI of Casp3 expression; $A_{ii}$: percentage of SST interneurons that are ablated in each layer of V1 at 5, 10, and 15 DPI. *** $P < 0.001$, two-way ANOVA test. Scale bar: 100 µm.

B. Distribution of PV interneurons after ablation of SST interneurons at 15 DPI. *Left*: Representative images of PV-tdT$^+$ interneurons in V1 from ctrl mice or those with selective expression of Casp3 in SST interneurons. *Right*: Number of PV-tdT$^+$ interneurons in each layer of V1 from ctrl or Casp3-injected mice (n = 5 mice). $P$ = 0.92, two-way ANOVA test. Scale bar: 100 µm.
