## Supplementary material for "Reciprocal interaction between cortical SST and PV interneurons in top-down regulation of retinothalamic refinement": Fig. S4

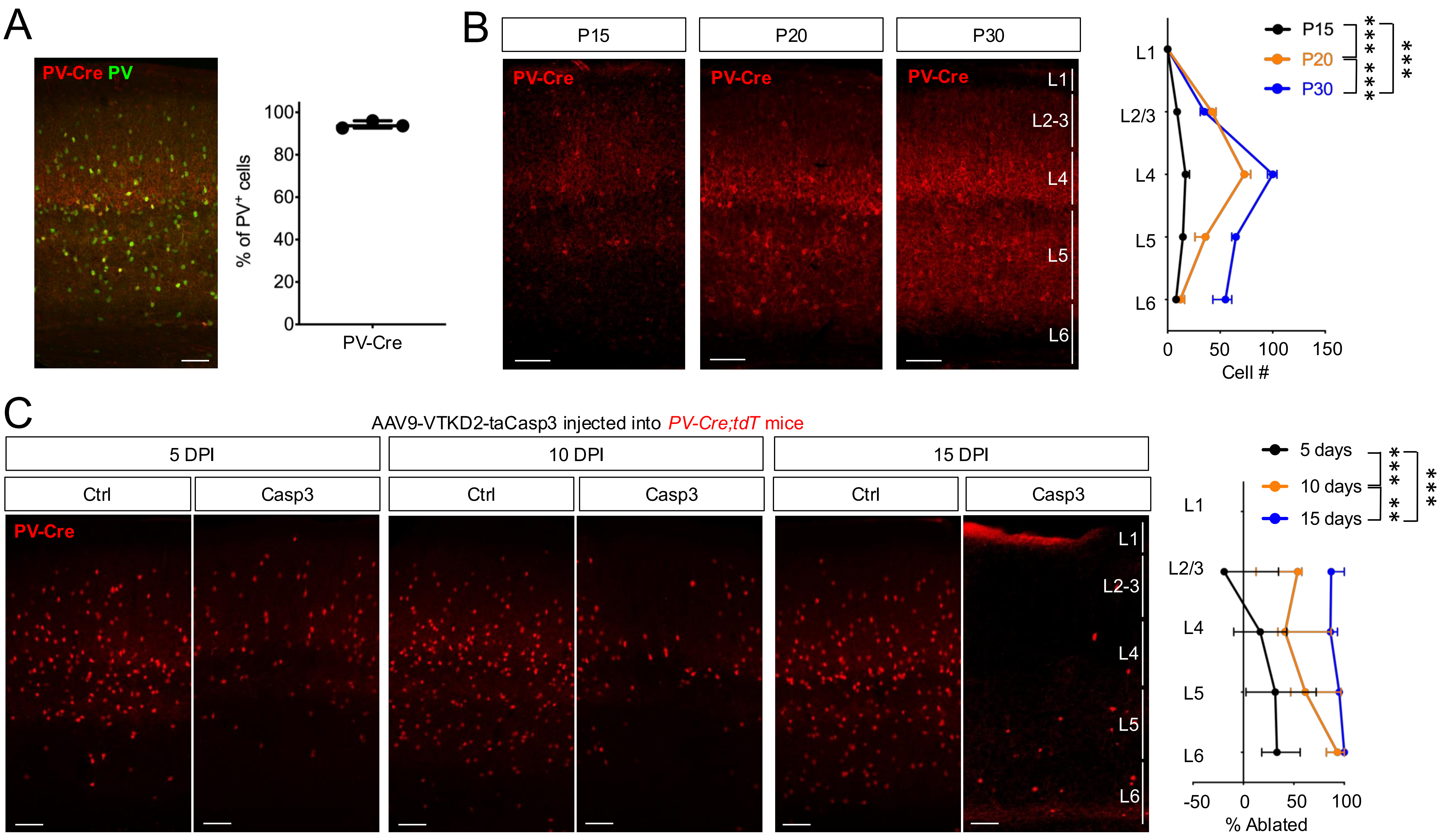


Figure S4. Ablation of PV interneurons in the V1 is detectable by 5 DPI, related to Figure 4.
A. Specificity of *Pvalb-Cre;tdT* transgenic mice in labeling PV$^+$ interneurons. *Left*: Confocal image of PV-Cre$^+$ neurons (red) from *Pvalb-Cre;tdT* mice co-stained with PV antibody (green). *Right*: Proportion of PV-Cre$^+$ interneurons that are labelled by PV antibody (n = 3 mice). Scale bar: 100 µm.
B. Developmental distribution of PV-Cre$^+$ interneurons from *Pvalb-Cre;tdT mice*. *Left*: Representative images of PV-Cre$^+$ interneurons at P15 (n = 4 mice), P20 (n = 3 mice), and P30 (n = 3 mice). *Right*: Developmental distribution of PV-Cre$^+$ interneurons across V1. *** $P < 0.001$, two-way ANOVA test. Scale bar: 100 µm.
C. Distribution of PV-Cre$^+$ interneurons with time after expression of Casp3 at DPI 5 (n = 3), 10 (n = 3), and 15 (n = 3). *Left*: Representative images of remaining PV-Cre$^+$ interneurons in the V1 after 5, 10, or 15 DPI of Casp3 expression; *Right*: Percentage of PV-Cre$^+$ interneurons that are ablated at 5, 10, and 15 DPI. ** $P < 0.01$, *** $P < 0.001$, two-way ANOVA test. Scale bar: 100 µm.
