## Supplementary material for "Reciprocal interaction between cortical SST and PV interneurons in top-down regulation of retinothalamic refinement": Fig. S5

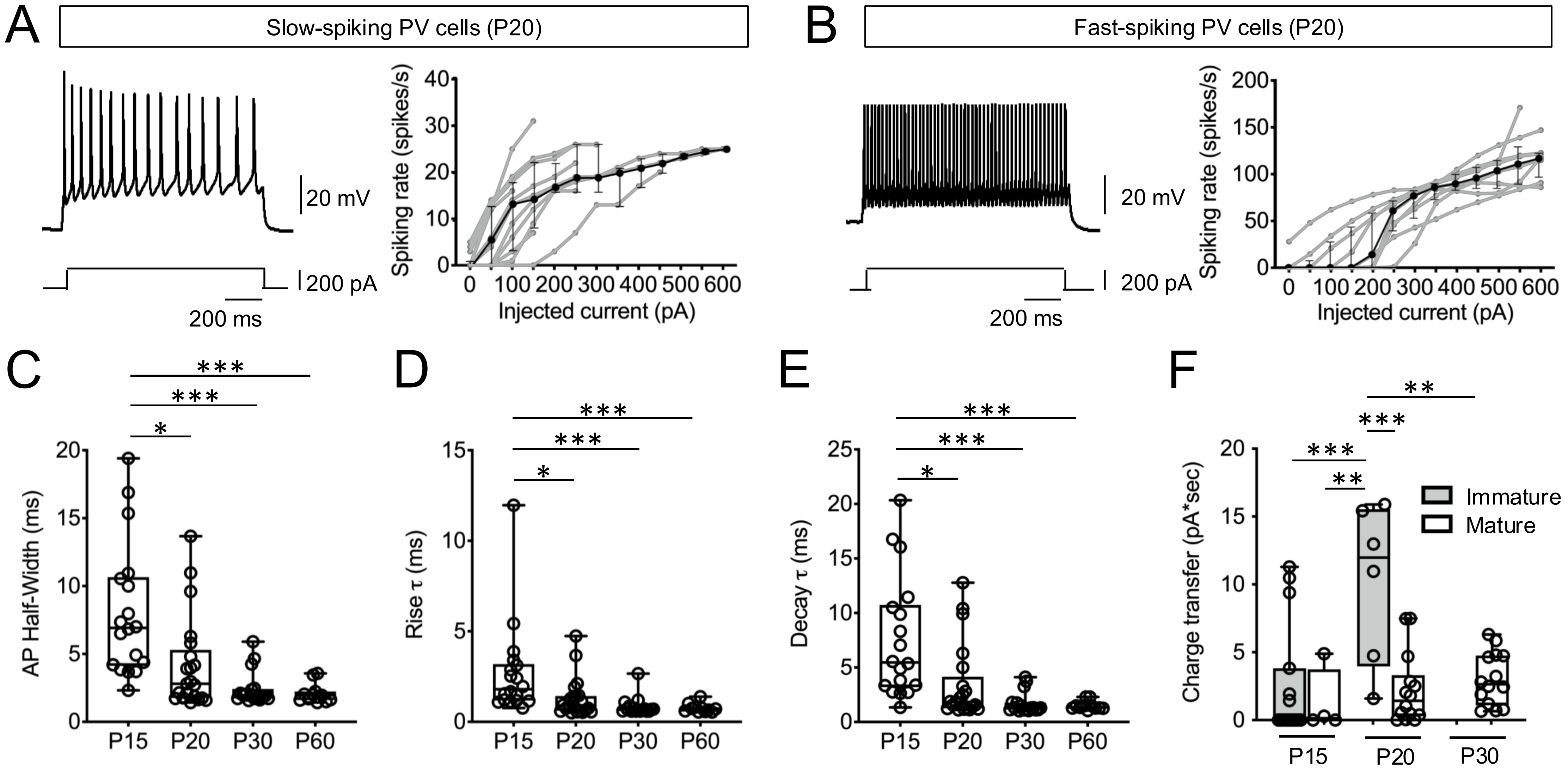


Figure S5. Firing properties of PV interneurons during development, related to Figure 6.
A. Firing pattern of slow-spiking PV interneurons. *left*: Representative recording from a slow-spiking PV interneuron in response to a current injection of 200 pA; *right*: Average input-output relationship of slow-spiking PV interneurons at P20. Grey lines: raw traces of input-output curve from each recording; Solid line: median input-output gain. n = 13 cells.
B. Firing pattern of fast-spiking PV interneurons. *left*: Spiking trace of fast-spiking PV interneuron with current injection of 200 pA; *right*: input-output gain of fast-spiking PV interneurons. n = 9 cells.
C. Half-width of action potentials of PV interneurons during development.
D. Rise τ of action potentials.
E. Decay τ of action potentials.
F. Charge transfer of SST→PV interneuron IPSCs sorted by immature vs mature PV interneurons.
C-F: * $P < 0.05$, ** $P < 0.01$, *** $P < 0.001$, two-way ANOVA, Sidak's multiple comparisons test.
