## Supplementary material for "Reciprocal interaction between cortical SST and PV interneurons in top-down regulation of retinothalamic refinement": Table S1

**SST→PN IPSCs (pA) during normal development**

| | P10 | P15 | P20 | P30 | P60 |
|---|---|---|---|---|---|
| n | 18 | 18 | 29 | 18 | 5 |
| median | 0 | 9.203 | 24.2 | 55.2 | 65.42 |
| IQR | 0-5.135 | 5.975-12.41 | 8.06-37.84 | 35.63-100 | 49.04-74.4 |

**SST→PN IPSCs (pA) after LDR**

| LDR | P30 | |
|---|---|---|
| | ctrl | LDR |
| n | 18 | 27 |
| median | 47.05 | 7.292 |
| IQR | 30.72-79.67 | 0-23.5 |

**SST→PN IPSCs (pA) after CDR**

| | P15 | | P30 | |
|---|---|---|---|---|
| | ctrl | CDR | ctrl | CDR |
| n | 8 | 18 | 15 | 27 |
| median | 7.075 | 9.581 | 35.19 | 23.84 |
| IQR | 4.375-20.68 | 5.573-14.73 | 18.13-47.05 | 11.88-65.47 |

**SST→PN IPSCs (pA) after ablation of SST interneurons**

| | P30 | |
|---|---|---|
| | ctrl | Casp3 |
| n | 21 | 24 |
| median | 48.06 | 0 |
| IQR | 28.48-71.77 | 0-13.11 |

Table S1. Amplitude of SST→PN IPSCs (pA). Related to Figures 1, 2, 3, and S2.
