## Supplementary material for "Reciprocal interaction between cortical SST and PV interneurons in top-down regulation of retinothalamic refinement": Table S2

| SST→PV IPSCs (pA) during normal development | | | | |
|---|---|---|---|---|
| | P15 | P20 | P30 | P60 |
| n | 19 | 20 | 14 | 13 |
| median | 0 | 110.8 | 108 | 143.1 |
| IQR | 0-162.2 | 56.25-291.4 | 64.78-177.6 | 104.5-164.5 |

| SST→PV IPSCs (pA) after LDR | | |
|---|---|---|
| | P30 | |
| | ctrl | LDR |
| n | 12 | 24 |
| median | 108 | 162.5 |
| IQR | 69.54-152.7 | 77.86-289.5 |

| SST→PV IPSCs (pA) after ablation of SST interneurons | | |
|---|---|---|
| | P30 | |
| | ctrl | Casp3 |
| n | 13 | 13 |
| median | 120.3 | 0 |
| IQR | 65.2-206.4 | 0-0 |

| SST→PV IPSCs (pA) during normal development | | | | | | |
|---|---|---|---|---|---|---|
| | P15 | | P20 | | P30 | |
| | Immature | Mature | Immature | Mature | Immature | Mature |
| n | 15 | 4 | 6 | 14 | 0 | 14 |
| median | 0 | 15.86 | 359 | 69.38 | 0 | 108 |
| IQR | 0-162.2 | 0-147.2 | 179.1-496 | 39.57-196.2 | 0 | 64.78-177.6 |

| SST→PV IPSCs (pA) after LDR | | | | |
|---|---|---|---|---|
| | P30 | | | |
| | ctrl | | LDR | |
| | Immature | Mature | Immature | Mature |
| n | 0 | 12 | 4 | 20 |
| median | 0 | 108 | 179.7 | 150.4 |
| IQR | 0 | 69.54-152.7 | 144.2-361.3 | 70.43-289.5 |

Table S2. Amplitude of SST→PV IPSCs (pA). Related to Figures 1, 2, and 3.
