## Supplementary material for "Reciprocal interaction between cortical SST and PV interneurons in top-down regulation of retinothalamic refinement": Table S3

**SST interneurons treated with Casp3 (ctrl: n = 39; Casp3: n = 44)**

| | Max EPSC | | SF EPSC | | FF | |
|---|---|---|---|---|---|---|
| | ctrl | Casp3 | ctrl | Casp3 | ctrl | Casp3 |
| median | 4.006 | 2.559 | 0.2136 | 0.04799 | 0.06018 | 0.0238 |
| IQR | 2.363-5.914 | 1.517-4.315 | 0.07438-0.4704 | 0.03188-0.08188 | 0.0316-0.1088 | 0.01304-0.08959 |

**PV interneurons treated with Casp3 (ctrl: n = 44; Casp3: n = 47)**

| | Max EPSC | | SF EPSC | | FF | |
|---|---|---|---|---|---|---|
| | ctrl | Casp3 | ctrl | Casp3 | ctrl | Casp3 |
| median | 3.39 | 3.546 | 0.1275 | 0.2038 | 0.04766 | 0.07275 |
| IQR | 2.259-4.095 | 2.454-4.725 | 0.06588-0.2689 | 0.07875-0.4363 | 0.0218-0.09523 | 0.03894-0.1411 |

**LDR+SST activation (Normal ctrl: n = 25; LDR ctrl: n = 25; LDR+Gq: n = 42)**

| | Max EPSC | | | SF EPSC | | | FF | | |
|---|---|---|---|---|---|---|---|---|---|
| | Normal ctrl | LDR ctrl | LDR+ Gq | Normal ctrl | LDR ctrl | LDR+ Gq | Normal ctrl | LDR ctrl | LDR+ Gq |
| median | 2.013 | 1.612 | 2.502 | 0.3029 | 0.0538 | 0.2167 | 0.1562 | 0.0444 | 0.0858 |
| IQR | 1.143-2.781 | 0.928-2.09 | 1.44-3.38 | 0.139-0.555 | 0.0378-0.163 | 0.110-0.449 | 0.049-0.343 | 0.0262-0.118 | 0.0381-0.238 |

**LDR+PV ablation (Normal ctrl: n = 21; LDR ctrl: n = 26; LDR+Casp3: n = 30)**

| | Max EPSC | | | SF EPSC | | | FF | | |
|---|---|---|---|---|---|---|---|---|---|
| | Normal ctrl | LDR ctrl | LDR+ Casp3 | Normal ctrl | LDR ctrl | LDR+ Casp3 | Normal ctrl | LDR ctrl | LDR+ Casp3 |
| median | 2.873 | 1.458 | 2.651 | 0.3256 | 0.05313 | 0.1741 | 0.1396 | 0.06332 | 0.1059 |
| IQR | 1.568-3.776 | 0.710-2.19 | 1.608-3.763 | 0.154-0.751 | 0.0323-0.171 | 0.111-0.75 | 0.0490-0.262 | 0.026-0.108 | 0.0371-0.275 |

Table S3. Measurement of retinogeniculate responses. Related to Figures 3, 4, and 5.
